# Triggered Release from Thermosensitive Liposomes Improves Tumor Targeting of Vinorelbine

**DOI:** 10.1101/2022.11.02.514937

**Authors:** Maximilian Regenold, Kan Kaneko, Xuehan Wang, H. Benson Peng, James C. Evans, Pauric Bannigan, Christine Allen

**Author notes:** **Corresponding author:** Christine Allen, PhD, Leslie Dan Faculty of Pharmacy, University of Toronto 144 College Street, Toronto, Ontario, M5S 3M2, Canada.

## Abstract

Triggered drug delivery strategies have been shown to enhance drug accumulation at target diseased sites in comparison to administration of free drug. In particular, many studies have demonstrated improved targetability of chemotherapeutics when delivered via thermosensitive liposomes. However, most studies continue to focus on encapsulating doxorubicin while many other drugs would benefit from this targeted and localized delivery approach. The proposed study explores the therapeutic potential of a thermosensitive liposome formulation of the commonly used chemotherapy drug vinorelbine in combination with mild hyperthermia (39-43 °C) in a murine model of rhabdomyosarcoma. Rhabdomyosarcoma, the most common soft tissue sarcoma in children, is largely treated using conventional chemotherapy which is associated with significant adverse long-term sequelae. In this study, mild hyperthermia was pursued as a non-invasive, non-toxic means to improve the efficacy and safety profiles of vinorelbine. Thorough assessment of the pharmacokinetics, biodistribution, efficacy and toxicity of vinorelbine administered in the thermosensitive liposome formulation was compared to administration in a traditional, non-thermosensitive liposome formulation. This study shows the potential of an advanced formulation technology in combination with mild hyperthermia as a means to target an untargeted therapeutic agent and result in a significant improvement in its therapeutic index.

**Graphical abstract:** 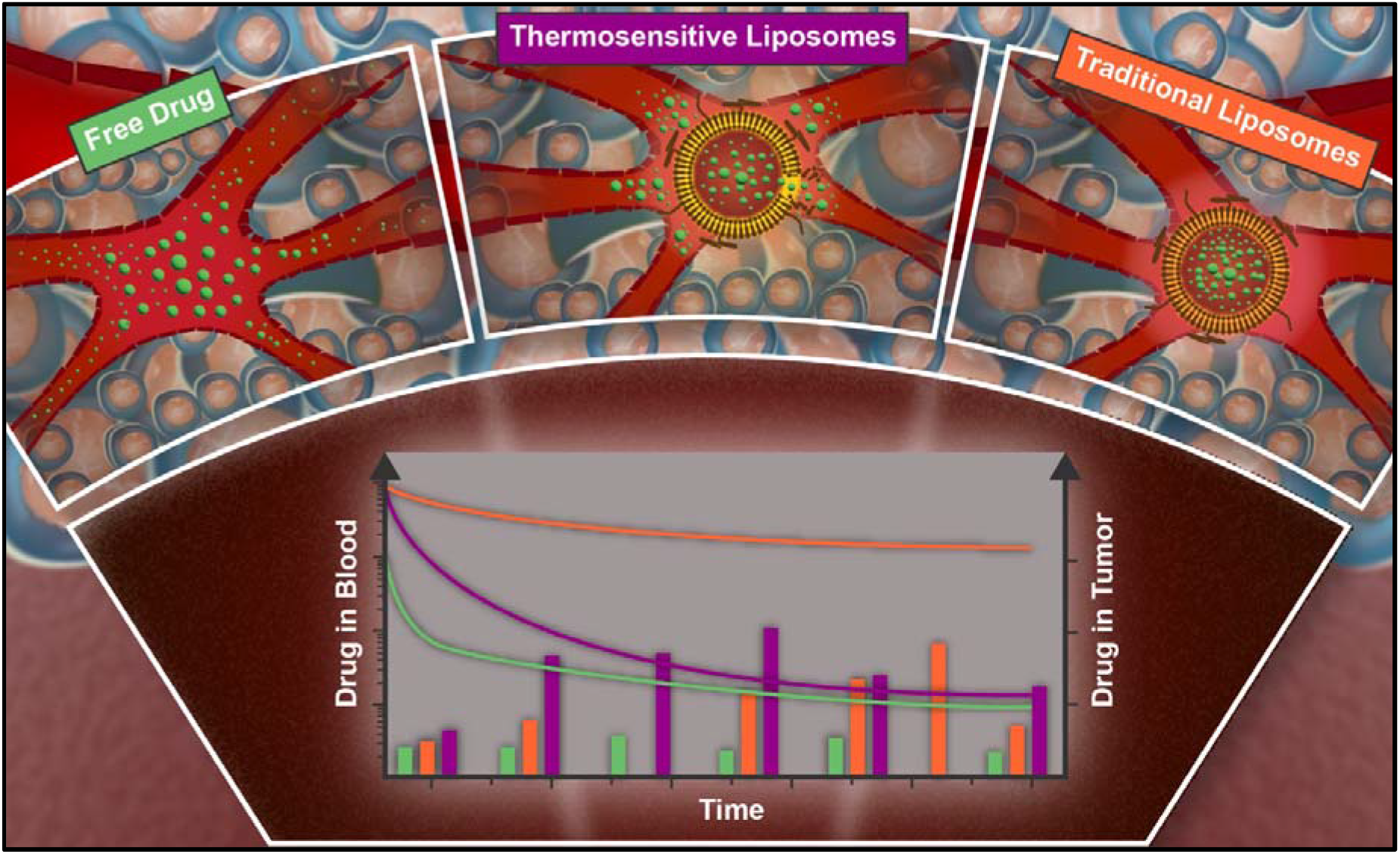

## 1. Introduction

Childhood cancer remains the leading cause of death by disease among Canadian children. Approximately 6 % of all childhood cancers are classified as soft tissue sarcomas (STSs) with rhabdomyosarcoma (RMS) being the most prevalent subtype (accounting for nearly 50 % of all STSs) [1]. This means that only 3-4 % of childhood cancer patients are diagnosed with RMS making it a relatively rare cancer. The treatment of childhood cancers in general has seen significant improvements over the course of the last 50 years. Large study groups, such as the Children’s Oncology Group (COG), have conducted comprehensive multicentre clinical trials and identified multimodal therapeutic protocols that resulted in significant increases in the overall five-year survival rates of pediatric oncology patients [2,3]. These studies have significantly shaped the treatment of RMS. In general, the treatment of RMS is guided by a series of risk factors and is normally comprised of chemotherapy, surgery, and radiotherapy. This treatment combination usually achieves survival rates between 70-90% for patients presenting with localized disease [4], However, approximately one-third of patients experience disease recurrence with the overwhelming majority (70-80 %) of recurrences localized at the primary disease site [5,6].

The backbone of RMS chemotherapy, in North America, remains a two- or three-drug combination comprised of intravenous administration of the conventional chemotherapeutic agents vincristine, actinomycin D, and/or cyclophosphamide (VAC). Yet the treatment of relapsed RMS remains a significant challenge with no universal standard treatment regimen identified to date [6]. Post-relapse chemotherapy regimens range from more aggressive and extensive VAC cycles to regimens including doxorubicin, cyclophosphamide, etoposide, and ifosfamide, and many others [6]. However, none of these drugs specifically target RMS or cancer cells in general and thus contribute to and/or result in significant toxicity.

Driven by a better understanding of disease mechanisms underlying RMS and with a goal towards improving outcomes and mitigating adverse effects in relapsed RMS, several clinical trials are assessing targeted therapy approaches [7,8]. For example, bevacizumab as well as temsirolimus in combination with vinorelbine (VRL) and cyclophosphamide were recently studied in a Phase II clinical trial where the addition of temsirolimus led to an improvement in event free survival [9]. Other drugs under investigation include PARP-(NCT03155620), HDAC-(NCT04308330), or ATR-inhibitors (NCT05071209) as well as immune checkpoint inhibition therapies (NCT02304458). However, the multitude of signaling pathways that are involved, as well as the rarity of the disease, pose significant challenges to the identification and development of these targeted therapy approaches [10]. As a result, current treatment options remain heavily reliant on conventional chemotherapy.

This vulnerable patient population is particularly at risk of experiencing and suffering from immediate and long-term adverse effects that are directly related to systemic exposure to conventional chemotherapy [11–13]. Additionally, the treatment of relapsed RMS often calls for even more aggressive chemotherapy regimens leading to significant systemic toxicities [14], Thus, drug delivery approaches that assure a more targeted deposition of drug, improve local control and reduce systemic toxicities may greatly benefit this patient population. A common method to achieve this, is the use of nanotechnology-based drug delivery approaches, specifically, liposomal encapsulation of chemotherapy drugs. In the current study, we explored a thermosensitive liposome delivery strategy that affords externally triggered release of the chemotherapy drug vinorelbine specifically at the heated tumor site.

Vinorelbine is a vinca-alkaloid that has previously been shown to be active in the treatment of recurrent RMS [15,16]. More recently, its promising activity in RMS was supported by encouraging results achieved when added as continued maintenance therapy in combination with low-dose cyclophosphamide [17]. The results of this study marked the first improvement in RMS survival rates from an experimental chemotherapy regimen in 30 years and have since led to a change in the European standard-of-care for treatment of recurrent RMS. However, it also raised several questions, particularly around which factor or combination of factors (e.g., prolonged chemotherapy treatment, efficacy of VRL and/or low-dose cyclophosphamide) led to the marked improvements [18]. These results in combination with previous studies hint towards an important role for VRL in future treatment of RMS [9]. Accordingly, VRL is currently being investigated as part of the induction chemotherapy drug combination (with actinomycin D and cyclophosphamide versus VAC) in a clinical trial conducted by the COG [NCT04994132].

Several research groups have encapsulated VRL in liposomes in an effort to increase drug delivery to the target disease site resulting in an improvement in local treatment control with a reduction in systemic toxicity [19–21], While this approach has shown promising results in the past (e.g., increased drug delivery to tumor and reduced systemic toxicity), traditional liposome-based drug delivery also faces several challenges. Namely, heterogeneity in the uptake and distribution of drug at tumour sites, due to variability in the enhanced permeability and retention (EPR) effect, and limited drug release from the liposomes [22,23]. To overcome these challenges thermosensitive liposomes were developed and pursued in combination with localized heating at the tumor site. Thermosensitive liposomes afford rapid drug release once heated to mild hyperthermic temperatures (39-43 °C) in the tumor vasculature. This creates a local concentration gradient of drug that drives the drug molecules into the surrounding tumor tissue [24], Thus, unlike drug delivery in traditional liposomes, this approach does not rely on long-circulating, nanocarriers that accumulate passively at the tumor site via the EPR effect. If designed properly, upon heating to mild hyperthermic temperatures, thermosensitive liposomes release near to 100% of their cargo enabling the drug to enter the tumor tissue as free molecules.

We previously developed and characterized a thermosensitive liposome formulation encapsulating VRL (ThermoVRL) [25]. In the current study, we assessed the *in vivo* performance of ThermoVRL in comparison to administration of free drug or drug encapsulated in a traditional, non-thermosensitive liposome formulation (NTSL-VRL) [20]. The results demonstrate that the liposome formulations of drug result in significant improvements in efficacy relative to that achieved with administration of free drug. ThermoVRL was found to deliver drug more specifically to the tumor in comparison to NTSL-VRL. Thus, this study demonstrates that heat-triggered release from thermosensitive liposomes can confer targetability to conventional, commonly used untargeted chemotherapy drugs.

## 2. Materials and Methods

### 2.1. Materials

Vinorelbine tartrate (VRL) and vinblastine sulfate (VB) were obtained from Selleck Chemicals (Houston, TX, USA). Sodium sucrose octasulfate (Na_8_SOS) was purchased from Toronto Research Chemicals (North York, ON, Canada). 1,2-Dipalmitoyl-sn-glycero-3-phosphocholine (DPPC), N-(carbonyl-methoxypolyethylenglycol 2000)-1,2-distearoyl-sn-glycero-3-phosphoethanolamine (PEG_2k_-DSPE), 1-stearoyl-2-lyso-sn-glycero-3-phosphocholine (lyso-SPC, MSPC) and cholesterol were obtained from Corden Pharma (Plankstadt, Germany). 1,2-Distearoyl-sn-glycero-3-phosphocholine (DSPC) was purchased from NOF Corporation (White Plains, NY, USA). Triethylamine (TEA), Dowex^®^ 50WX8-200 and 50WX4 200-400, bovine serum albumin (BSA, heat shock fraction, pH 7, ≥98 %), fetal bovine serum (FBS), penicillin and streptomycin (P/S), 2-mercaptoethanol and Hoechst 33342 were purchased from Sigma-Aldrich (Oakville, ON, Canada). Sepharose CL-4B was purchased from GE Healthcare Bio-Sciences (Mississauga, ON, Canada). Rh30 cells were provided by the Children’s Oncology Group Childhood Cancer Repository. Cell culture medium and insulin-transferrin-selenium supplement (ITS) were purchased from Life Technologies (Burlington, ON, Canada).

### 2.2. Cell culture and evaluation of *in vitro* cytotoxicity

The human RMS cell line Rh30 was cultured in Iscove’s Modified Dulbecco’s Medium supplemented with 20 % FBS, 1 % P/S, 4 mM L-Glutamine and 1× ITS. Cells were kept at 37 °C with 5 % CO_2_ unless otherwise indicated. Cells were seeded in 96-well plates at 1500 cells/well for 24 h followed by treatment with VRL for 1 h. Cells were then washed and incubated for a total of 72 h. Cytotoxicity was determined via the acid phosphatase assay: aliquots of 2 mg/mL phosphatase substrate p-nitrophenylphosphate were added to cells for 1 h followed by the addition of 0.1 N NaOH and UV absorbance measurement at 405 nm. Half maximal inhibitory concentration (IC_50_) values were obtained using GraphPad Prism 6.0 (GraphPad Software, San Diego, CA, USA) to fit the data with a 4-parameter sigmoidal dose response curve. The effect of mild hyperthermia (HT) on drug cytotoxicity was determined through exposing cells to a temperature of 42 °C for the first hour of the total 72 h incubation period.

### 2.3. Liposome preparation

Both non-thermosensitive liposomes (NTSL-VRL) and thermosensitive liposomes (ThermoVRL) were gradient loaded with VRL using triethylammonium sucrose octasulfate (TEA_8_SOS). NTSL-VRL was prepared following a protocol previously developed by Drummond et al. [20] and ThermoVRL was prepared as described in our previously published manuscript [25]. In brief, DSPC, cholesterol, and PEG_2k_-DSPE at a molar ratio of 3/2/0.015 (i.e., 59.8/39.9/0.3) were solubilized in 65 % (v/v) ethanol at 60-65 °C and then combined with 10 volumes of pre-heated TEA_8_SOS solution (0.65 M sulfate group concentration) to achieve a lipid concentration of 125 mM. ThermoVRL was prepared by dissolving lipids in chloroform and drying them to produce a lipid film composed of DPPC, MSPC, and PEG_2k_-DSPE at a molar ratio of 86/10/4. The film was subsequently hydrated in TEA_8_SOS solution (0.22 M-0.65 M sulfate group concentration) at approximately 55 °C for a minimum of 30 min. Both liposome formulations were extruded (Lipex Extruder, Northern Lipids, Vancouver, BC, Canada) three times through double stacked track-etch polycarbonate membranes (Whatman Inc., Clifton, NJ, USA) with a 200 nm pore size and 10 times through 100 nm pore sized membranes at 200 and 400 psi nitrogen pressure, respectively. NTSL-VRL liposomes were extruded at 70 °C and ThermoVRL liposomes at 55 °C. Unencapsulated TEA_8_SOS was removed by dialysis overnight (4 °C; 50 kDa MWCO) in a 1000-fold volume excess of HEPES-buffered saline (HBS; 20 mM HEPES, 150 mM sodium chloride, pH 6.5). To load NTSL-VRL with VRL, a 10 mg/mL solution of VRL tartrate in water was added to liposomes at 350gVRL/mol lipid, the pH was adjusted to 6.5 using 1 N NaOH followed by incubation at 60 °C for 30 min. ThermoVRL liposomes were similarly loaded either at 80 or 30 g VRL/mol lipid for one hour at 35 °C. Liposomes were subsequently chilled on ice for 10 minutes and unencapsulated drug was removed by another over-night cycle of dialysis in HBS. ThermoVRL liposomes were concentrated via tangential flow filtration using a polysulfone MicroKros^®^ filter (Spectrum, Rancho Dominguez, CA, USA). All formulations (including VRL solubilized at 3 mg/mL in 0.9 % saline solution; free-VRL) were filtered through a 0.22 μm PES filter into a sterile rubber-sealed glass vial (ALK-Abelló Inc., Round Rock, TX, USA) prior to characterization or use in animal experiments.

### 2.4. Liposome characterization and evaluation of *in vitro* drug release

Liposome size was measured using dynamic light scattering (DLS) (Zetasizer Nano ZS, Malvern Instruments, Malvern, WOR, UK) and an intensity-based analysis after a 1: 100 dilution in phosphate buffered saline pH 7.4 (PBS). The zeta potential was measured using the same instrument following a 1: 100 dilution in Milli Q water. The melting phase transition temperature (T_m_) was determined using a TA Q100 differential scanning calorimeter (TA Instruments, New Castle, DE, USA) where samples were subjected to three heating cycles between 25 °C and 60 °C at a rate of 1 °C/min. VRL concentrations were measured using an HPLC-MS method previously described [25].

VRL release from NTSL-VRL was determined by adding 200 μL of liposomes to 10 mL pre-heated PBS with 45 g/L BSA at 37 and 42 °C. These solutions were stirred in an incubator at the respective temperatures for the duration of the experiment. At specific timepoints, 200 μL aliquots were removed and released drug was separated from the liposomes via size exclusion chromatography with Sephadex CL-4B gel columns. Drug levels were then quantified using the aforementioned HPLC-MS method.

### 2.5. Tumor model and localized mild hyperthermia treatment

All animal studies were conducted in accordance with the guidelines of the Animal Care Committee of the University Health Network (UHN, Toronto, ON, Canada). Aliquots of 1.5 × 10^6^ Rh30 cells in OptiMEM/Matrigel (1/1) (Corning^®^ Matrigel^®^ Basement Membrane Matrix, LDEV-free) were injected subcutaneously into the right flank of 5-6-week-old female SCID mice to develop ectopic xenograft tumors. Once palpable lesions were detectable, tumor growth was tracked with caliper measurements of width (*w*) and length (*l*). The following equation was then used to calculated tumor volumes: 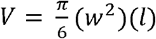. Treatment was started once tumor volumes reached >90 mm^3^, and was administered on days 0, 7, and 14 for tumor sizes < 12 mm in the longest dimension. Tumors were heated to 42.5 °C for 5 min prior to intravenous tail-vein administration followed by another 20 min of heating. A laser-based heating system previously developed and described by Dou et al. was used to achieve localized and conformal heating [26]. In brief, a 763 nm diode laser (Model CD 403 laser, Ceralas, Jena, Germany) connected to a custom-built illuminator (Spectralon, Labsphere Inc., North Sutton, NH, USA) via a 400 μm fiber was used to deliver a homogenous light distribution (± 15%). The power was manually adjusted between 0.2 to 0.65 W/cm^2^ to maintain a target temperature of 42.5 °C. A centrally placed point-based optical fiber temperature probe (Luxtron Model 790, LumaSense Technologies Inc., Santa Clara, CA, USA) within the tumor was used to monitor the temperature. Tumor dimensions of > 15 mm in any dimension or a body weight loss > 20 % were selected as ethical endpoints.

Tumor growth rates were determined using the Gompertz model. In brief, the tumor growth data was fitted to the following equation using the Excel solver function: *V* (*t*) = *αe^-βe^-γt^^;* where V(t) is the tumor volume (in mm^3^) at time point t (in days), *a* and ß are constants, and γ is the volumetric growth rate (in mm^3^/day).

### 2.6. Evaluation of tumor vascularization

Mice with tumors > 90 mm^3^ were anesthetized with 5 % isoflurane and euthanized by cervical dislocation followed by rapid tumor resection. Tumors were embedded in OCT compound (Tissue-Tek, Sakura Finetek, Torrance, CA, USA), flash-frozen in liquid nitrogen, and stored at −80 °C. Sections of 5 μm thickness were cut from the frozen block, stained and imaged for blood vessels (with anti-CD31 IF staining; 560/607 nm) in combination with DAPI (358/461 nm). A fluorescence scanner (AxioScan Zeiss), located at The Advanced Optical Microscopy Facility (AOMF, Toronto, ON, Canada) was used to image whole-mount tissue sections at 0.5 μm/pixel. Tissue preparation, staining and imaging were performed by the correlative pathology laboratory at the STTARR Innovation Centre (STTARR, Toronto, ON, Canada). Images were processed using the HALO^®^ Image Analysis Platform (Indica Labs Inc., Albuquerque, NM, USA).

### 2.7. Evaluation of *in vivo* pharmacokinetics of VRL administered as free drug, in NTSL-VRL, or ThermoVRL combined with localized mild HT in tumor bearing mice

The pharmacokinetics profiles of VRL in tumor bearing mice (19.3 ± 1.4 g) were investigated after administration as free drug (i.e. free-VRL, n = 44), NTSL-VRL (n = 40), or ThermoVRL (n = 44) at 15 mg/kg in combination with mild HT (as described above; Section 2.5). 250 μL gastight glass syringes (1725 LT SYR, Hamilton Company, Reno, NV, USA) connected to an intravenous tail-vein catheter were used to ensure accurate administration of VRL (as free-VRL, NTSL-VRL, or ThermoVRL). Animals receiving free-VRL or ThermoVRL plus mild HT were sacrificed via cardiac puncture while anesthetized (5 % isoflurane) at 1, 3, 5, 15, 20, 30, 60, 120, 180, and 360 min post administration. Mice receiving NTSL-VRL plus mild HT were sacrificed at 1, 5, 20, 60, 120, 180, 360, 720, and 1440 min post administration. The heating and sampling process is outlined in Figure 7. Additionally, animals receiving free-VRL, NTSL-VRL or ThermoVRL without the addition of mild HT were sacrificed at 20 min post administration. This timepoint was chosen to allow comparison with studies on formulations previously developed and studied by our research group [27], Blood was added to heparinized tubes and heart, lungs, liver, kidneys, spleen, and tumors were resected, washed in PBS and flash frozen in liquid nitrogen. Blood, frozen tissues, and tumors were stored at −80 °C prior to analysis. 10 % tissue homogenates in Milli Q were prepared using a Precellys 24-Dual (Bertin Instruments, Montigny-le-Bretonneux, France) tissue homogenizer. Stock solutions of VRL and VB were prepared in MeOH at 1 mg/mL and stored at −20 °C. Working solutions of VRL were prepared by diluting the VRL stock solution with MeOH/H_2_O (1/1; v/v). Standards with final concentrations between 0.5 to 1000 ng/mL were obtained by spiking and mixing 45 μL of blank whole blood or tissue homogenate with 5 μL of the working solutions.

**Figure 1:**
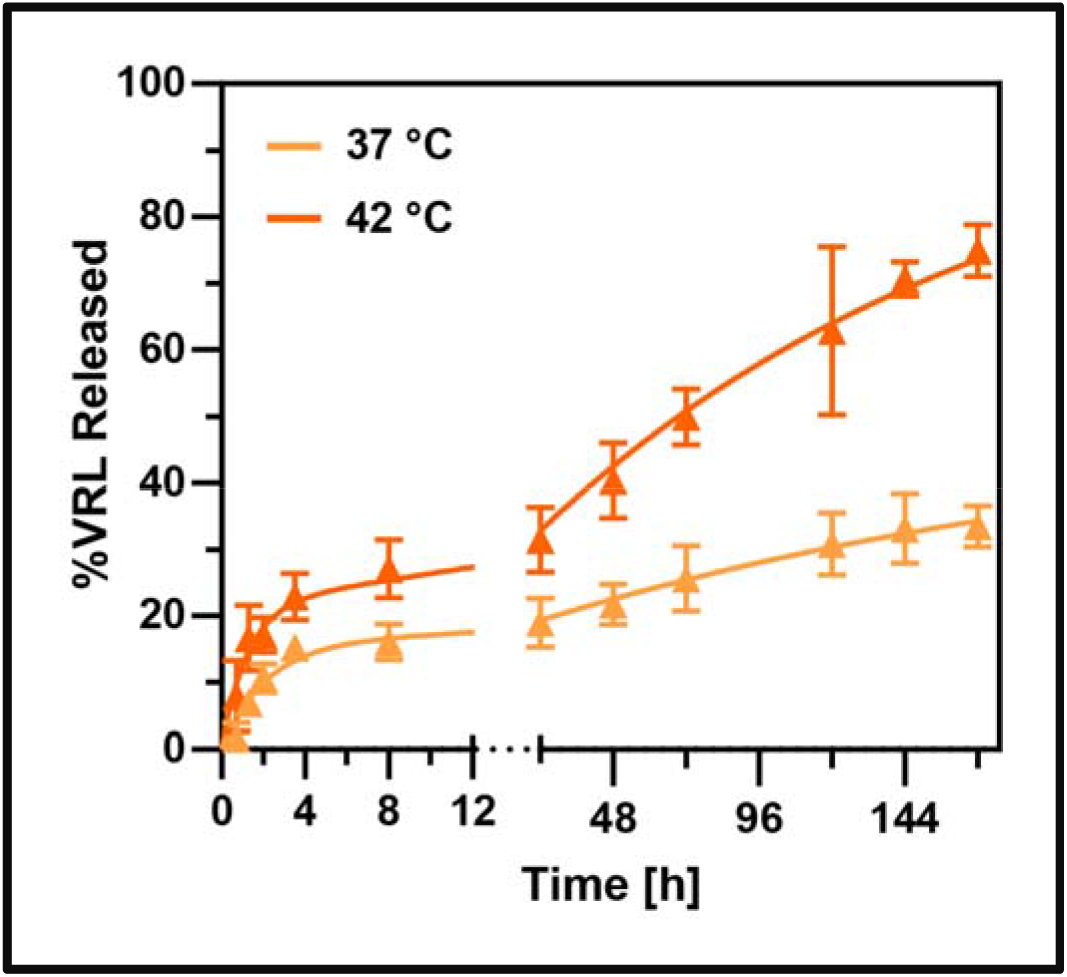
*In vitro* release of VRL from non-thermosensitive liposomes at 37 and 42 °C in release media containing 45 g/L BSA. Statistically significant differences in drug release at the two temperatures were only detected at time points greater than 1.3 h. Error bars represent SD (n = 3).

**Figure 2:**
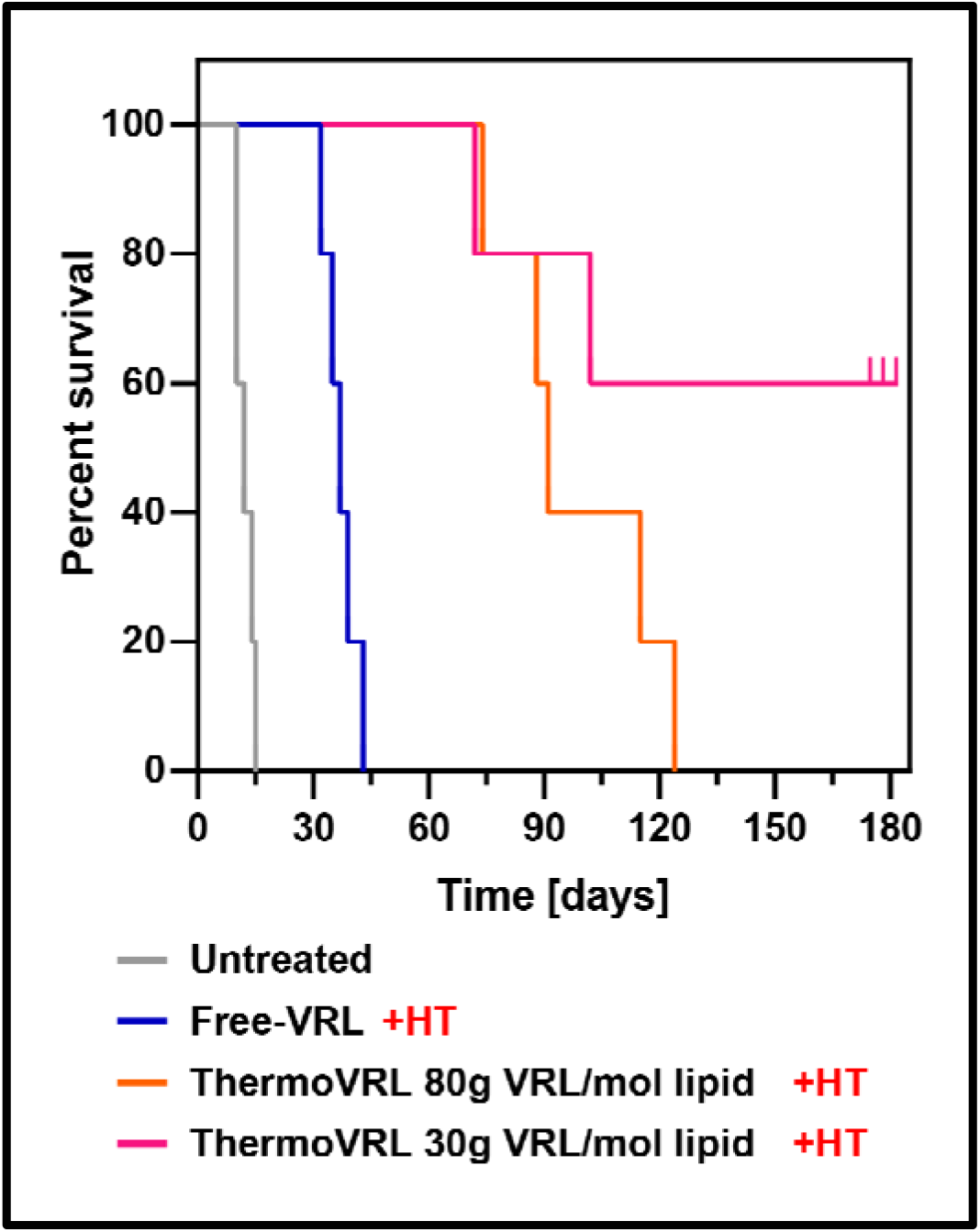
Kaplan-Meier survival analysis of tumor bearin female SCID mice treated with free vinorelbine, o thermosensitive liposomes loaded at different drug-to-lipii ratios in combination with mild hyperthermia (42.5 °C 25 min) localized to the tumor site. Treatments wer administered at a dose of 15 mg/kg on days 0, 7, and 14 (n = 5).

**Figure 3:**
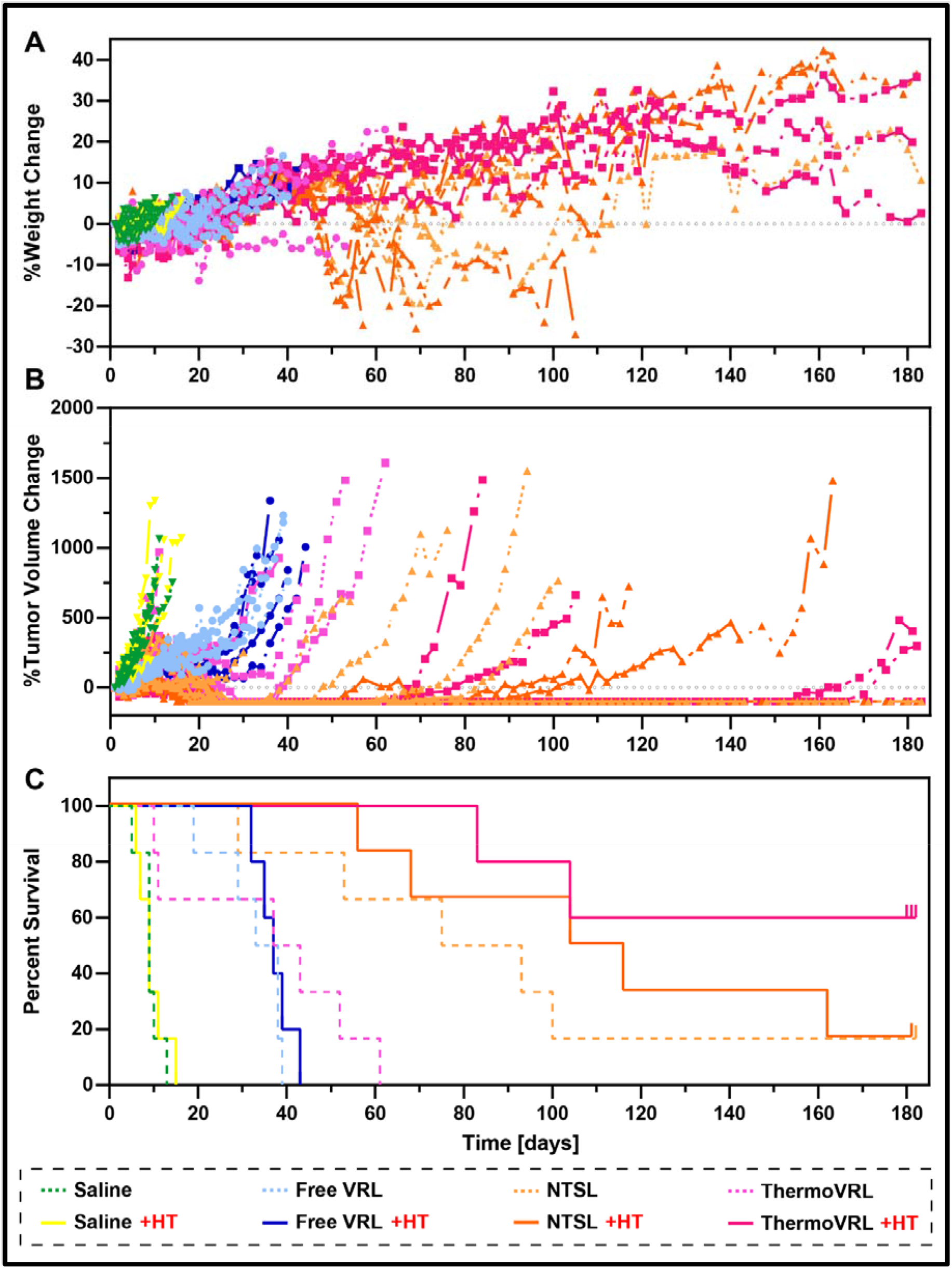
Efficacy study comparing control and treatment groups including mice receiving saline, free-VRL, non-thermosensitive liposomal VRL (NTSL-VRL), and thermosensitive liposomal VRL (ThermoVRL)) with and without the addition of mild hyperthermia localized to the tumor site. Female SCID mice bearing subcutaneous Rh30 xenografts were treated at 15 mg/kg on days 0,7, and 14. A) Percent body weight change. B) Percent tumor volume change and C) Kaplan-Meier survival analysis. ThermoVRL + HT and NTSL-VRL + HT significantly increased the median survival time compared to administration of free-VRL(p <.05). (n ≥ 5).

**Figure 4:**
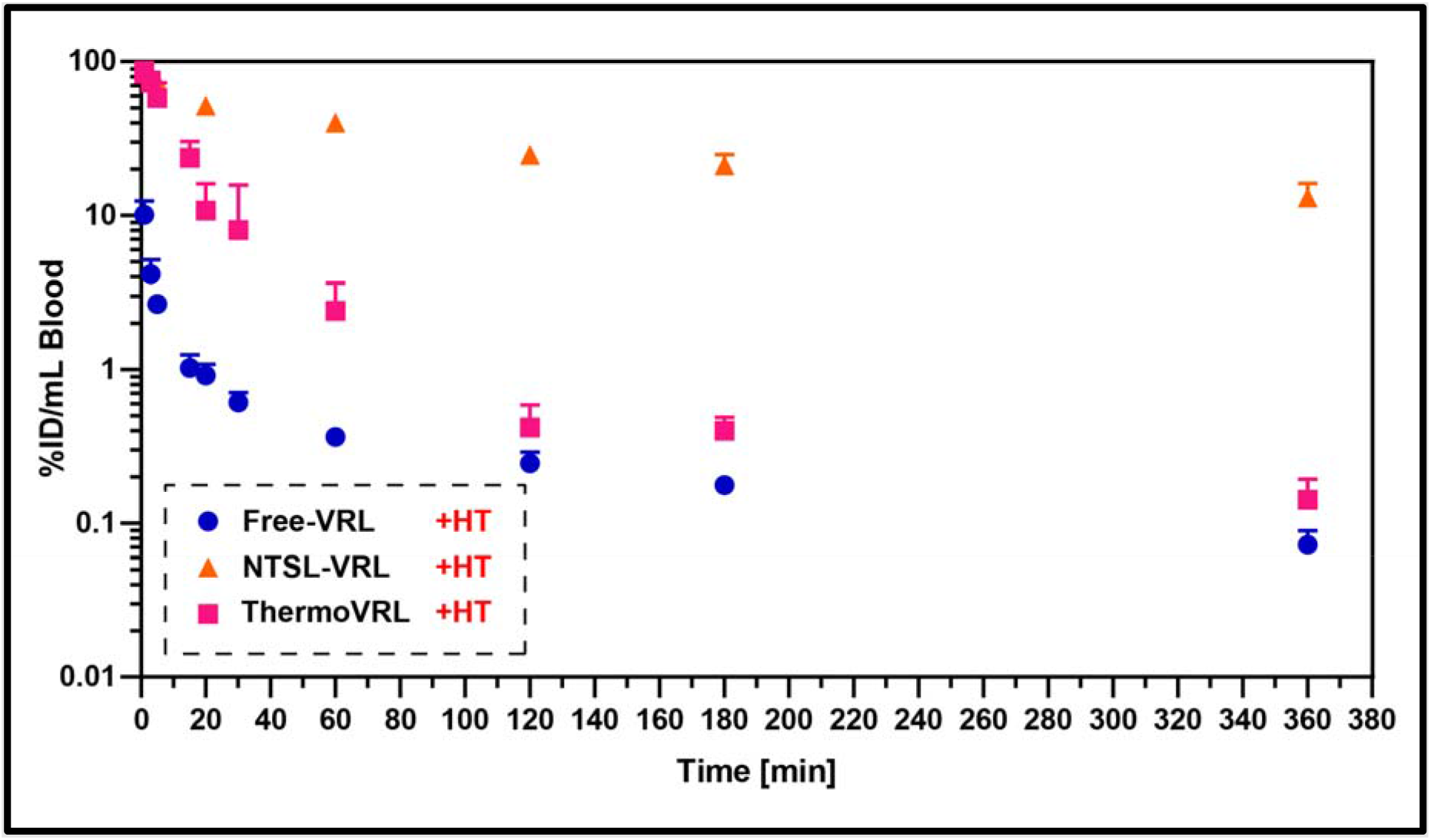
Concentration-time profiles for vinorelbine (VRL) administered as free-VRL, NTSL-VRL, or ThermoVRL, in combination with mild hyperthermia localized to the tumor (42.5 °C, 25 min), in whole blood following tail-vein injection. Data is presented as mean ± SD (n = 4).

**Figure 5:**
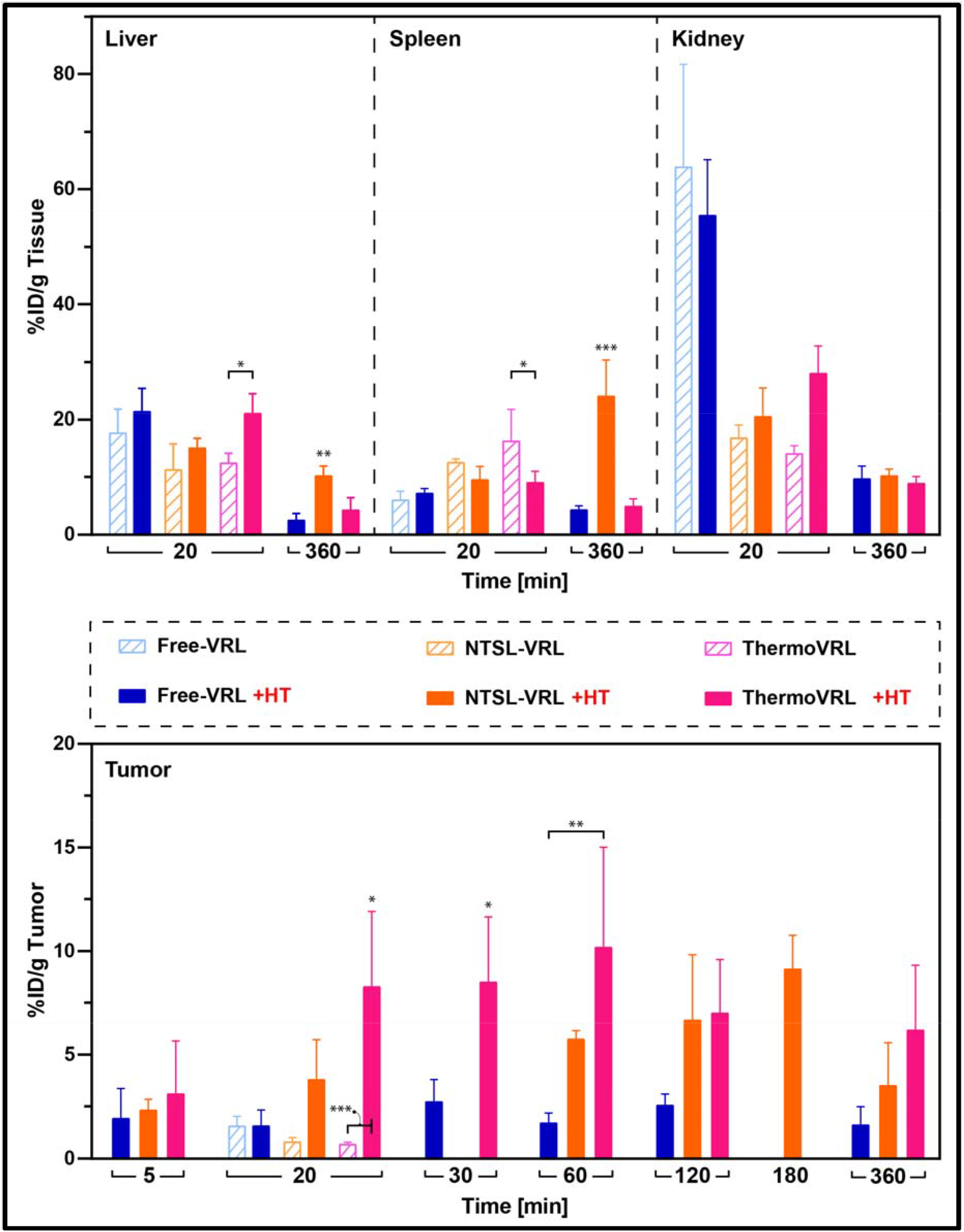
Biodistribution and tumor distribution of VRL at various timepoints post administration of free-VRL, NTSL-VRL, or ThermoVRL. VRL distribution in liver, spleen, and kidneys was assessed at an early (20 min) and late (360 min) timepoint. Tumor distribution of VRL was evaluated in mice treated with free-VRL and ThermoVRL after 5, 20, 30, 60, 120, and 360 min and after 5, 20, 60, 120, 180, and 360 min for NTSL-VRL. The distribution after 20 min is shown for treatments administered without (dashed bars) and with mild HT localized to the tumor (filled bars). Statistical significance of measurements within each timepoint (i.e., 5 min, 20 min, etc.) was determined by one-way ANOVA with Bonferroni correction and data is presented as mean ± SD (n = 4).

**Figure 6:**
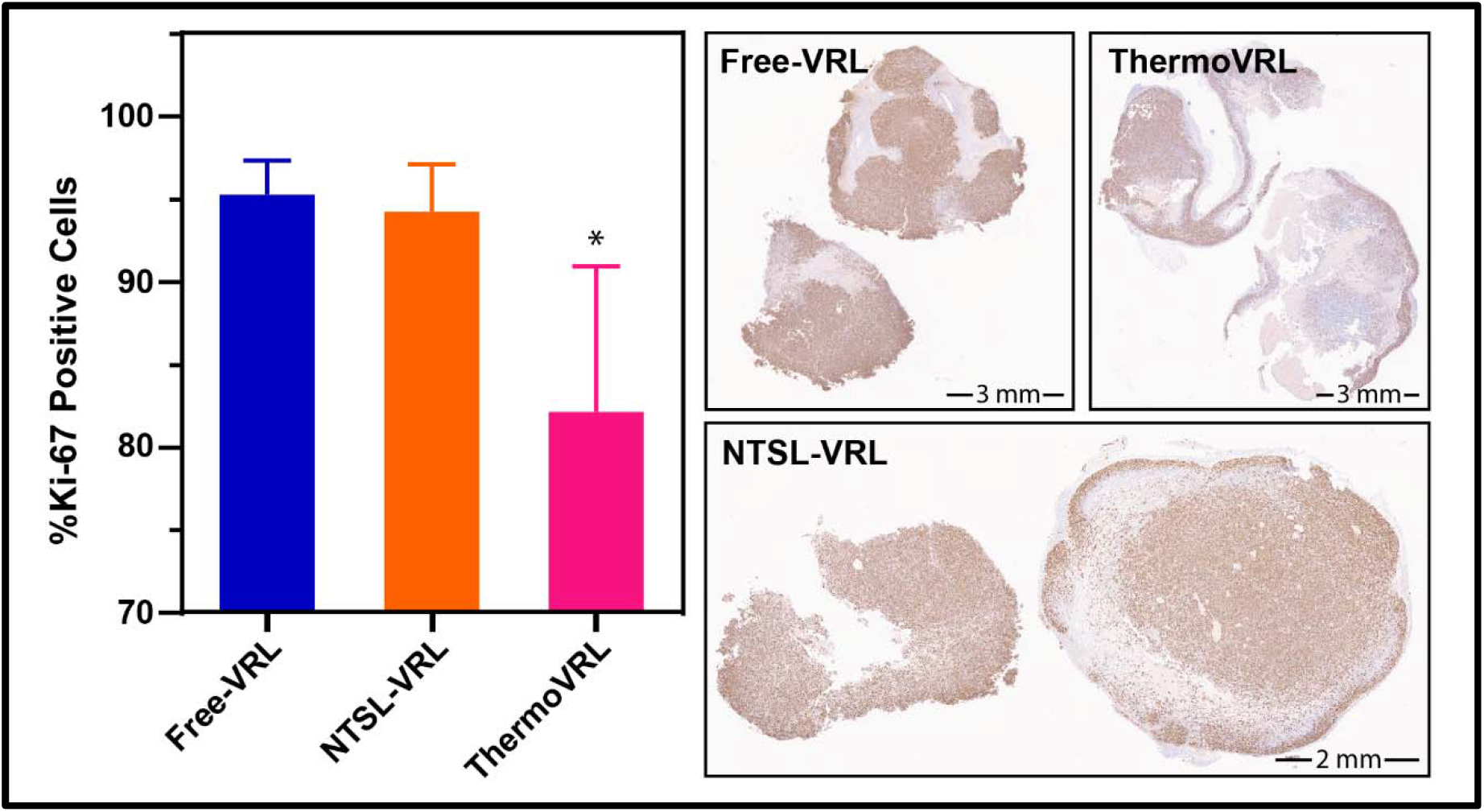
Rh30 tumor sections stained for Ki-67 24 h post treatment with free-VRL, NTSL-VRL, or ThermoVRL at 15 mg/kg in combination with mild hyperthermia localized at the tumor site. %Ki-67 cell positivity is significantly reduced in tumor sections of animals treated with thermosensitive liposomal VRL compared to treatment with free VRL or traditional liposomal VRL.

**Figure 7:**
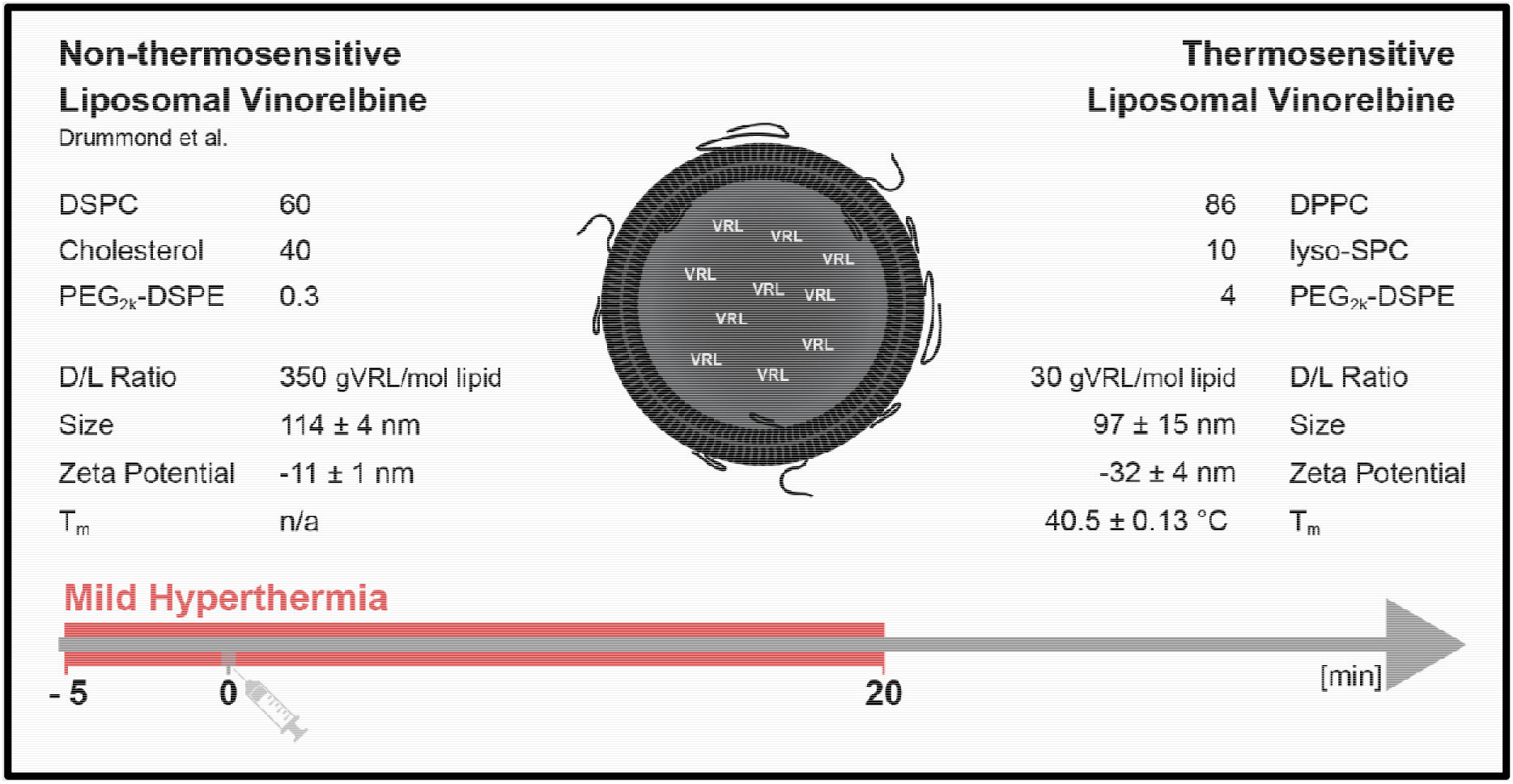
Physico-chemical properties of non-thermosensitive liposomal VRL (composition reported previously by Drummond et al. [20]) and thermosensitive liposomal VRL (ThermoVRL) [25]. The liposome formulations are actively loaded with VRL using a TEA_8_SOS gradient followed by removal of unencapsulated drug by dialysis. Size was determined by DLS and the melting phase transition temperature (T_m_) by DSC. Values reported here are in agreement with our previously published study [25]. Data is presented as mean ± SD of at least three independent liposome batches.

VRL was extracted by adding 950 μL of MeOH/ACN (1/1; v/v) containing 35 ng/mL VB to 50 μL whole blood or tissue homogenate. Samples were vigorously vortex-mixed for 3 min and centrifuged at 14,000 rpm for 10 min at 4 °C (Eppendorf 5804 R). The clear supernatant was then transferred to a separate vial, dried using nitrogen gas at 35 °C, resuspended in ACN/H_2_0 (1/1; v/v) and centrifuged prior to transferring into an autosampler glass vial. VRL concentrations were subsequently analyzed by HPLC-MS.

Blood, tissue, and tumor concentrations of VRL delivered as free-VRL, NTSL-VRL, and ThermoVRL were normalized to the dose and expressed as %ID/ml or %ID/g tissue, respectively. Normalization of sample concentration to the dose facilitated data comparison even if there were slight incongruities in the dose administered. Noncompartmental analysis by data regression was conducted for blood, tissue, and tumor data. The area under the curve to time infinity (AUC_∞,blood_) for blood was determined by the trapezoidal rule, which is the summation of the area up to the last sampling point (AUC_0-last,blood_) and extrapolated area after the last sampling point (C_last,blood_) divided by the terminal decay rate constant. The total (blood) body clearance (CL_blood_) after i.v. dosing via tail-vein injection was determined by dosei.v,/AUC_∞blood_. Tissue and tumor partitioning coefficients (K_p,tissue:blood_) were obtained by dividing tissue or tumor over blood AUC from 20 to 360 min (AUC_20-360 min,tissue_/AUC_20-360 min,blood_).

### 2.8. Analysis of blood chemistry and tissue histology

Tumor bearing mice were treated with free-VRL, NTSL-VRL or ThermoVRL at 15 mg/kg via tail vein injection in combination with mild HT. Animals were euthanized after 24 h and 7 days by cardiac puncture under 5 % isoflurane anesthesia and blood was collected in potassium EDTA-tubes for a complete blood count (CBC) analysis using the VetScan_®_ HM5 Hematology System (Abaxis, Union City, CA, USA). Additionally, blood was collected in lithium heparinized tubes for analysis using the VetScan VS2 Chemistry Analyzer in combination with the VetScan Prep Profile II reagent rotor. Tissues (heart, lungs, liver, kidneys, and spleen) and tumors were excised, fixed with 10 % neutral buffered formalin for 24-48 h before transfer into 70 % EtOH followed by embedding in paraffin. 5 μm thick serial sections were cut and stained with H&E to assess cellular morphology of the tissue. Immunohistochemistry with a Ki-67 antibody (Invitrogen PA5-19462) was performed to stain for proliferating cells on the tissue and a TUNEL assay was conducted to demonstrate cell death caused by the treatment. As mentioned previously, images of the H&E and IHC stained sections were processed using the HALO^®^ Image Analysis Platform.

### 2.9. Statistical analysis

Statistical analysis was performed using SSPS Statistics 28.0 (IBM, Armonk, NY, USA). One-way ANOVA with Bonferroni post hoc testing was performed to evaluate the *in vitro* characterization parameters (e.g., size, zeta potential, T_m_) between ThermoVRL loaded with VRL at 30 or 80 g VRL/mol lipid and NTSL-VRL. *In vitro* drug release at specific time points from NTSL-VRL at different temperatures was compared by unpaired t-test. Differences in mouse survival were calculated and compared by log-rank test and Bonferroni correction for multiple comparisons. Differences in body weight changes were determined by unpaired t-test between two groups at specific timepoints. Statistical significance of *in vivo* parameters (e.g., t_1/2_, AUC_∞blood_, CL_blood_, K_p,tissue:blood_) between free-VRL, NTSL-VRL, and ThermoVRL at specific timepoints were analyzed by one-way ANOVA with Bonferroni post hoc testing.

## 3 Results

### 3.1. Liposome characterization

ThermoVRL liposomes were loaded with either 30 or 80 g VRL/mol lipid and different concentrations of the internal trapping agent TEA_8_SOS, 0.22 M and 0.65 M, respectively. NTSL-VRL liposomes were prepared at 350 g VRL/mol lipid with 0.65 M internal TEA_8_SOS concentration [20]. NTSL-VRL liposomes were marginally larger in size (114 ± 4nm, p < .01) and had a slightly less negative zeta potential (− 11 ± 1 mV, p < .001) than both ThermoVRL liposome formulations. These differences in size and surface charge are statistically significant, however, it is not expected that they will influence their *in vivo* behaviour. As previously reported, the melting phase transition temperature (T_m_) of the ThermoVRL formulation loaded with less drug (30 g VRL/mol lipid) was found to be slightly higher compared to the same formulation loaded with more VRL (80 g VRL/mol lipid) (40.5 ± 0.13 °C and 39.8 ± 0.05 °C; p < .001). Thermosensitive liposomes are known to release their encapsulated content once heated to their T_m_. Thus, the slightly higher T_m_ observed for the formulation with the lower drug loading level points towards improved stability of this formulation at lower temperatures [24].

Non-thermosensitive liposomes were loaded at an initial D/L ratio of 350 g VRL/mol lipid, with drug loaded liposomes exhibiting a size of 114 ± 4 nm and a zeta potential of −11 ± 1 mV. As shown in **Figure 1,** the *in vitro* release data for the NTSL-VRL formulation was fitted with a first order equation comprised of two parts: one accounting for the initial fast drug release and another modeling the subsequent slower drug release phase (R(t)=R_max_(1-A e^-k1t^-(1-A) e^-k2t^)) **(Table 1**). VRL release was calculated to be 2 and 5 % after 20 min at 37 and 42 °C, respectively, and statistically significant differences in release at the different temperatures were only detected for time points after 1.3 h (p < .045). The *in vitro* release experiments demonstrate the significantly increased stability of NTSL-VRL compared to ThermoVRL (i.e. maximum release of almost 100 % achieved after 30 s at 42 °C; published previously in [25]) as well as the negligible effect of heating on drug release from NTSL-VRL, over relevant *in vivo* heating durations (here 20 min).

**Table 1:**
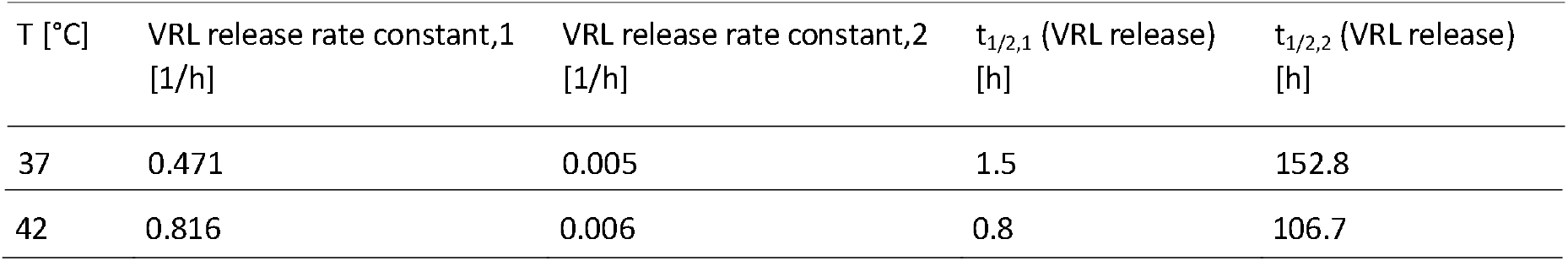
*In vitro* VRL release parameters for the non-thermosensitive liposome formulation of VRL (NTSL-VRL).

As shown in **Figure 2,** treatment with free-VRL + HT at 15 mg VRL/kg body weight significantly reduced the tumor growth rate compared to untreated mice (p < .001) leading to a significant increase in median survival time (MST) (37 ± 2 versus 12 ± 2 days; p < .02). MST of mice treated with ThermoVRL (at 15 mg VRL/kg weight) loaded with 30gVRL/mol lipid + HT (MST ≥ 180 days; p < .02) or 80 g VRL/mol lipid + HT (MST = 91 ± 3 days, p < .02) was significantly increased compared to untreated mice or mice receiving free-VRL + HT. No statistically significant difference (p = .676) in MST of animals treated with ThermoVRL loaded with different D/L ratios was observed. However, treatment with ThermoVRL loaded with 30 g VRL/mol lipid led to complete remission in three out of five mice, while none of the mice receiving ThermoVRL loaded with 80 g VRL/mol lipid survived past day 124. No statistically significant differences in body weight change were found between animals receiving ThermoVRL compared to animals receiving free-VRL (p > .09). Animals recovered rapidly following an initial body weight drop immediately after treatment compared to untreated control animals and body weight changes returned to baseline prior to second or third treatments.

Moving forward, only ThermoVRL liposomes loaded with 30 g VRL/mol lipid were considered for further *in vivo* evaluation. As shown in **Figure 3,** comparison between animals treated with saline or saline + HT did not result in statistically significant differences in tumor growth rates between groups (p = 1.0). As well, there was no difference in tumor growth rates between groups receiving free-VRL with or without the addition of mild HT (p = .35). While the addition of mild HT to ThermoVRL treatment did result in a significant increase in MST (37 ± 20 vs. > 180 days; p < .04), adding mild HT to the treatment with NTSL-VRL did not result in any statistically significant differences (111 ±7 vs. 75 ±24 days; p = 1.0). ThermoVRL + HT and NTSL-VRL + HT increased MST nearly five- and three-fold respectively, compared to free-VRL+HT (37 ± 2 days). Administration of ThermoVRL+HT resulted in complete tumor remission until day 150 post-treatment in three out of five animals followed by tumor recurrence in two out of the three mice while only one out of six mice treated with NTSL-VRL ± HT remained in complete remission for the duration of the study (180 days). Two out of six mice treated with NTSL-VRL+HT were euthanized due to body weight loss > 20 % and mice receiving NTSL-VRL ± HT experienced significantly greater body weight loss compared to mice treated with ThermoVRL + HT (when greatest body weight loss was observed for animals receiving NTSL-VRL ± HT; p < .05).

### 3.2. Tumor blood microvessel density and blood perfusion

Immunostaining for CD31 determined a blood microvessel density of 4.2 ± 1.9 % area within untreated tumors (> 90 mm^3^, 5 independent tumor xenografts). For comparison, using a similar approach Banerjee et al. found a comparable microvessel density of 4.86 ± 0.58 % area in the well vascularized MDA-MB-231 tumor model (ectopic subcutaneous implantation) [28,29].

### 3.3. Pharmacokinetics of VRL administered as free-VRL, NTSL-VRL, or ThermoVRL in combination with localized mild hyperthermia

The pharmacokinetic profiles for VRL following administration in different formulations (free-VRL, NTSL-VRL, or ThermoVRL) are provided in **Figure 4.** Clear differences in the VRL concentration-time profiles are observed. Data regression analysis determined VRL C_0_ (in blood) to be approximately 100 % dose/mL for NTSL-VRL and ThermoVRL, and only 12 %dose/mL when administered as free-VRL. Free VRL is rapidly cleared from blood and thus early sampling timepoints are crucial to accurately determining VRL C_0_ values [20]. However, the experimental setup of this study did not allow for blood sampling at earlier timepoints than one minute post treatment administration. The apparent half-life (t_1/2_), estimated by terminal phase regression of the log-linear portions of the blood decay curves were 141 ± 33, 291 ± 122, and 176 ± 127 min for free-VRL, NTSL-VRL, and ThermoVRL, respectively, although not statistically different (p > .212) (see **Table 2).** However, there were significant differences determined for the extrapolated AUC_∞blood_ normalized to the dose for free-VRL (189 ± 92 %dose*min/mL), NTSL-VRL (15465 ± 3788 %dose*min/mL), and ThermoVRL (1331 ± 155 %dose*min/mL), which provided different clearance values of 30.4 ± 10.5 mL/min/kg, 0.3 ± 0.1 mL/min/kg, and 4.1 ± 0.5 mL/min/kg, respectively. As expected, encapsulation in NTSL-VRL significantly reduced VRL clearance compared to administration as free drug (100-fold difference; p < .001). Similarly, although to a lesser extent (8-fold difference), VRL encapsulation in ThermoVRL reduced its clearance compared to free-VRL (p < .001). Lysolipid-containing thermosensitive liposomes are known to be significantly less stable *in vivo* compared to other thermosensitive liposomal formulations as well as traditional liposomes [30,31]. In addition, as expected, the volume of distribution determined from %dose/C_0_ was highest for free-VRL (9.97 ± 5.02 mL) and significantly reduced for NTSL-VRL (1.17 ± 0.11 mL) and ThermoVRL (1.03 ±0.32 mL).

**Table 2:** Noncompartmental parameters for vinorelbine (VRL) following intravenous administration as free-VRL, non-thermosensitive liposomal VRL (NTSL-VRL), and thermosensitive liposomal VRL (ThermoVRL), with each treatment in combination with mild hyperthermia localized to the tumor.

| Parameters | Units | Free-VRL + HT | NTSL-VRL + HT | ThermoVRL + HT |
| --- | --- | --- | --- | --- |
| Mouse weight | [g] | 19.9 ± 1.1 | 19.3 ± 1.7 | 18.7 ± 1.4 |
| AUC<sub>0-∞, blood</sub> | [%Dose*min/mL] | 189 ± 92 | 15465 ± 3788 | 1331 ± 155 |
| $t_{1/2, \text{terminal}}$ | [min] | 141 ± 33 | 291 ± 122 | 176 ± 127 |
| CL<sub>blood</sub> | [mL/min/kg] | 30.4 ± 10.5 | 0.3 ± 0.1 | 4.1 ± 0.5 |
| AUC<sub>20-360 min, liver</sub> | [%Dose*min/g liver] | 566 ± 177 | 1322 ± 150 | 766 ± 235 |
| AUC<sub>20-360 min, spleen</sub> | [%Dose*min/g spleen] | 594 ± 86 | 2386 ± 530 | 693 ± 176 |
| AUC<sub>20-360 min, kidney</sub> | [%Dose*min/g kidney] | 1843 ± 256 | 1477 ± 144 | 1406 ± 202 |
| AUC<sub>20-360 min, tumor</sub> | [%Dose*min/g tumor] | 746 ± 183 | 2219 ± 515 | 2550 ± 544 |
| $K_{p, \text{liver: blood}}$ | AUC<sub>liver</sub>/AUC<sub>blood</sub> | 4.43 ± 1.16 | 0.14 ± 0.01 | 0.60 ± 0.19 |
| $K_{p, \text{spleen: blood}}$ | AUC<sub>spleen</sub>/AUC<sub>blood</sub> | 4.71 ± 0.77 | 0.25 ± 0.05 | 0.55 ± 0.16 |
| $K_{p, \text{kidney: blood}}$ | AUC<sub>kidney</sub>/AUC<sub>blood</sub> | 14.56 ± 1.67 | 0.15 ± 0.01 | 1.10 ± 0.17 |
| $K_{p, \text{tumor: blood}}$ | AUC<sub>tumor</sub>/AUC<sub>blood</sub> | 5.93 ± 1.56 | 0.23 ± 0.04 | 2.01 ± 0.52 |

Upon closer examination, the tissue partition coefficients, K_P,tissue:blood_, determined for liver, spleen, and tumor for VRL administered as free-VRL were found to be between 4 to 6, with no statistically significant difference between these values (p = .8) **(Table 2** and **Figure 8).** However, the K_p,kidney:blood_ for kidney (i.e. 14.6 ±1.67) was significantly higher in comparison to values obtained for any of the other organs analyzed (p < .001). As expected, systemic i.v. administration of VRL does not provide tumor-selective tissue distribution. In general, the K_p_ ratios for VRL administered as NTSL-VRL were significantly lower compared to administration as free-VRL (p < .001). However, no statistically significant difference was found between spleen and tumor K_p_ ratios (0.25 ± 0.05 vs. 0.23 ± 0.04; p = 1.0) indicating a very similar VRL partitioning into both tissues when administered as NTSL-VRL. Comparable to the administration of free-VRL, this highlights that the delivery of VRL encapsulated in NTSL-VRL does not specifically target tumor tissue over other tissues **(Figure 8).**

**Figure 8:**
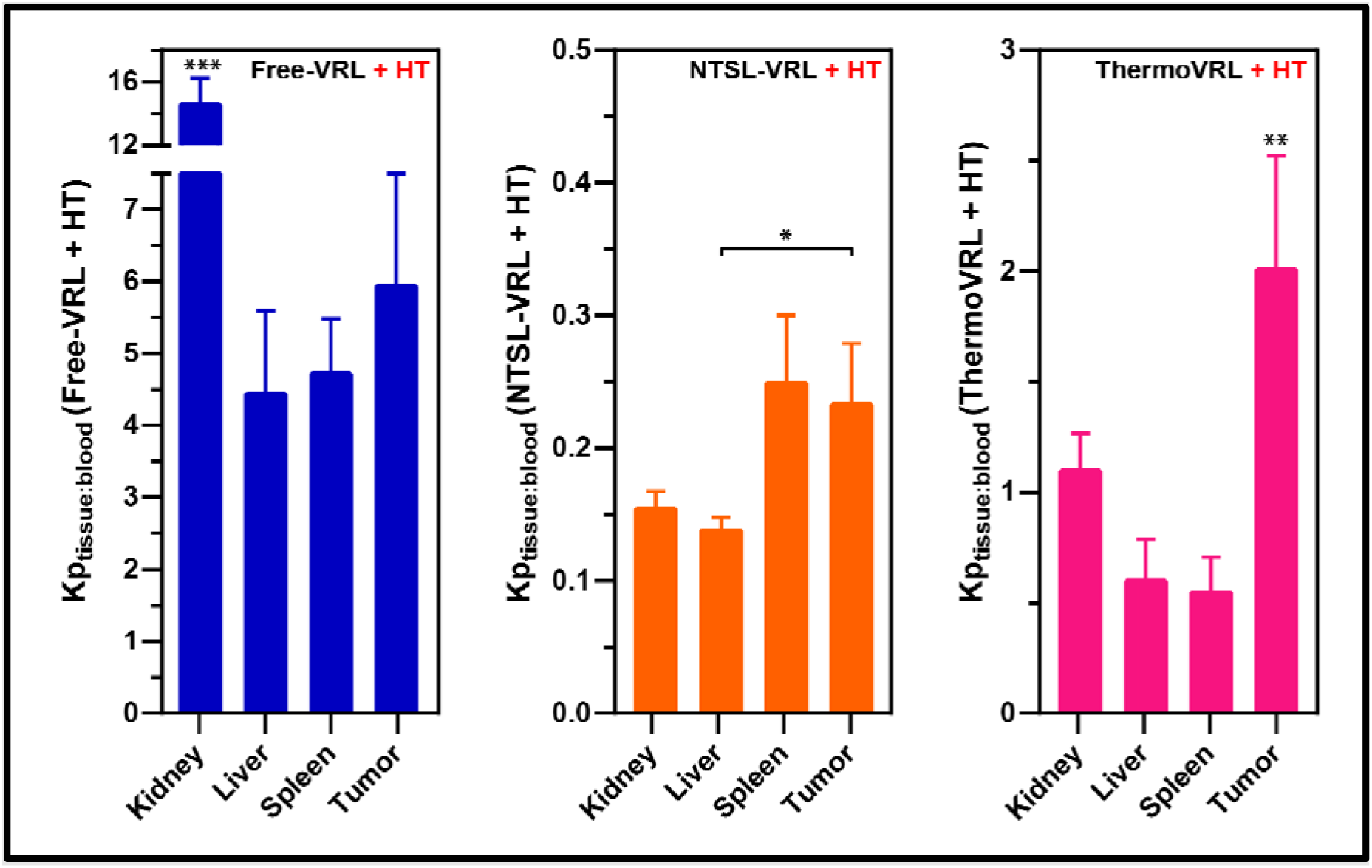
VRL distribution into tissues and tumor after administration of free-VRL, NTSL-VRL, or ThermoVRL at 15 mg VRL/kg body weight in combination with localized mild HT in tumor bearing female SCID mice. Data is presented as mean ±SD (n = 4). Treatment with ThermoVRL +HT provides tumorspecific delivery of VRL.

In contrast, administration of VRL as ThermoVRL provided the most intriguing results with K_p,tissue:blood_ values generally found to be between free-VRL and NTSL-VRL **(Table 2** and **Figure 8).** ThermoVRL + HT treatment led to a significantly higher tumor K_p_ ratio compared to K_P,tissue:blood_ ratios for any other tissue analyzed (p < .006). Specifically, the tumor K_p_ ratio was found to be more than three-fold higher compared to liver and spleen ratios and nearly two-fold higher than the kidney ratio. Moreover, the K_p,tumor:blood_ value for administration of ThermoVRL was calculated to be almost 10-fold higher than for NTSL-VRL. Many factors influence the partitioning of a specific drug from blood into the respective tissue, including its physico-chemical properties (e.g., lipophilicity, pKa, molecular weight) as well as transporter distribution or tissue composition [32]. This needs to be taken into consideration when comparing the partition coefficient of different drugs, but more importantly, different formulations of the same drug. A drug’s ‘effective’ physico-chemical properties are significantly altered when encapsulated in a liposomal formulation. Since free VRL is relatively lipophilic (logP of 4.65 (calculated) [25]) and membrane permeable, it is to be expected that liposomal encapsulation significantly reduces tissue uptake and thus K_p,tissue:blood_ values. Aside from the differences in VRL release (upon heating) from both liposomal formulations evaluated here, their different lipid compositions as well as drug retention influence VRL’s tissue partitioning. These are factors that need to be taken into account when comparing K_p,tissue:blood_ values obtained for NTSL-VRL+HT and ThermoVRL + HT. Altogether, this data shows that the administration of VRL via ThermoVRL in combination with localized mild HT is able to provide enhanced tumor specific drug delivery. In particular, in comparison to VRL administered as free drug (free-VRL + HT) or encapsulated in traditional liposomes (NTSL-VRL + HT).

It is important to note that pharmacokinetics of drugs administered in thermosensitive liposomes are commonly investigated in healthy animals [26,33–36]. Such non-tumor bearing experimental setups usually preclude analysis in combination with localized heating. This leads to the assessment of pharmacokinetic profiles of drugs encapsulated in triggered drug release systems without the actual external trigger applied to elicit drug release. However, as shown in **Figure S2,** localized heating dramatically affects the VRL concentration in whole blood when administered via ThermoVRL. In order to design optimized treatment protocols (i.e., duration and timing of heating as well as dosing of the TSL formulation) it is crucial to evaluate the drug’s pharmacokinetics in combination with triggering release.

### 3.4. Biodistribution of VRL following administration as free-VRL, NTSL-VRL, or ThermoVRL in combination with localized mild hyperthermia

The biodistribution of VRL after administration as free-VRL, NTSL-VRL, or ThermoVRL, in combination with mild HT, was assessed by measuring the concentration of drug in liver, kidney, and spleen both 20 and 360 min post administration **(Figure 5).** Additionally, biodistribution of VRL was determined 20 min post administration as ThermoVRL, NTSL-VRL, or free-VRL, without the addition of mild HT. It is important to note, that throughout the experiments presented in this manuscript the concentration of VRL and not the distribution of the delivery system (e.g. liposomes) was determined. VRL itself is mainly metabolized and extracted via the liver [37]. As shown in **Figure 5,** adding mild HT to the administration of ThermoVRL and thus triggering VRL release within tumor blood vessels increased liver VRL concentrations by nearly 60 % at 20 min post administration of drug (p < .04) while no statistically significant differences were observed for NTSL-VRL ± HT (p = 1.0) or free-VRL ± HT (p = 1.0). Wash-out of free VRL after triggered release from ThermoVRL within tumor blood vessels does increase the amount of systemically circulating free VRL explaining this significant increase in VRL concentrations in the liver after the addition of mild HT to ThermoVRL. However, as expected, a 2-fold increase in VRL concentration in the liver for NTSL+ HT treatment compared to ThermoVRL + HT or free-VRL + HT was detected 360 min post administration (10.2 ±1.7 versus 4.2 ± 2.2 and 2.5 ± 1.2 %ID/g liver; p < .003). Traditional liposomes are known to significantly prolong circulation lifetimes and to accumulate in organs of the mononuclear phagocyte system (MPS; e.g., liver and spleen) [38]. In accordance, the AUC_20-360 min,liver_ was found to be sig⋂ĩfĩcan11y higher for animals treated with NTSL+HT compared to ThermoVRL+HT or free-VRL+HT (p < .008), with no difference between ThermoVRL+HT or free-VRL ÷HT(p>.5) (Table 2).

No statistically significant difference in VRL concentrations in the spleen among groups + HT at 20 min post administration of drug were detected (p = .250). The comparison of treatment groups ± HT only found a statistically significant difference for ThermoVRL ± HT where the addition of mild HT led to a 55% decrease in VRL concentrations (p < .04) (Figure 5). At 360 min post administration, VRL concentrations in the spleen were increased 5-fold for NTSL-VRL + HT compared to both other groups + HT (23.9 ± 6.5 versus 4.8 ± 1.4 and 4.2 ± 0.8; p < .001). Similarly, the AUC_20-360 min,spleen_ for NTSL-VRL+HT treated animals was found to be significantly increased (p < .001) compared to ThermoVRL + HT or free-VRL + HT with no difference detected between the latter (p = 1.0) (Table 2).

As mentioned above, liposomes are known to preferentially accumulate in both the liver and spleen (as part of the MPS). As expected, NTSL-VRL led to significantly increased drug deposition in both organs. However, this off-target deposition of drug was remarkably reduced when VRL was administered as ThermoVRL in combination with mild HT. This provides evidence of efficient drug release from thermosensitive liposomes within tumor blood vessels prior to carrier accumulation in MPS organs.

Nearly 2-fold higher VRL concentrations in the kidney were measured for free-VRL ± HT compared to treatment with ThermoVRL ± HT or NTSL-VRL ± HT 20 min post administration (p < .005). The addition of mild HT did not significantly change kidney concentrations for any of the treatment groups ±HT (p > .534). Equally, no differences in the concentration of VRL in the kidneys at 360 min post administration between the different treatment groups + HT were found (p > .836). In consequence, the administration of VRL as free-VRL+HT led to a significantly increased AUC_20-360 min, kidney_ compared to administration as ThermoVRL + HT (p = .044), yet no statistically significant difference in AUC_20-360 min,kidney_ was found when administered as NTSL-VRL (p = .098).

### 3.5. Bulk tumor accumulation of VRL delivered via free-VRL, NTSL-VRL, or ThermoVRL in combination with localized mild HT

As illustrated in Figure 7, tumors were pre-heated for 5 min, followed by administration of free-VRL, NTSL-VRL, or ThermoVRL via tail-vein injection and another 20 min of heating post injection. Temperatures of heated tumors determined via a centrally placed single point temperature probe were 42.3 ±0.5 °C. No statistically significant difference in concentrations of VRL in the tumor were found between animals treated with ThermoVRL + HT, NTSL-VRL + HT or free-VRL + HT at 5, 120, and 360 min post administration (p > .053) (Figure 5). However, treatment with ThermoVRL + HT led to significantly higher VRL concentrations in the tumor 20 min post administration compared to NTSL ±HT or free-VRL ±HT (p < .026). Specifically, ThermoVRL +HT resulted in an over 2-fold increase in VRL concentration in the tumor compared to NTSL-VRL ! HT and an over 5-fold increase compared to free-VRL ± HT. Adding mild HT to free-VRL did not change VRL concentrations in the tumor (p = 1.0) and while the addition of mild HT to NTSL-VRL led to a nearly 5-fold increase in VRL concentration in the tumor, this difference was not found to be statistically significant (p = .38). However, combining ThermoVRL with mild HT led to more than a 12-fold increase in VRL concentrations in the tumor 20 min post administration compared to ThermoVRL alone (p < .001). VRL concentrations in the tumor at 30 min post administration were only assessed for animals treated with ThermoVRL +HT or free-VRL + HT, where a 3-fold increase in VRL concentration was detected for ThermoVRL + HT (p < .02). At 60 min post administration, no difference in VRL concentrations at the tumor was detected in animals treated with ThermoVRL +HT and NTSL-VRL + HT (p = .164); while, VRL concentrations in the tumor in animals treated with free-VRL + HT were significantly lower than that achieved with ThermoVRL + HT (p < .01). A similar pattern was revealed when the AUC_20-360 min,tumor_ for the different treatment groups + HT was calculated and compared **(Table 2).** While a larger AUC_20-360 min,tumor_ was obtained in animals treated with ThermoVRL + HT compared to NTSL-VRL + HT, this difference was not statistically significant (p = .964). And as expected, the estimated AUC_20-360 min,tumor_ for animals treated with free-VRL + HT was found to be significantly lower than the values obtained for animals treated with liposomal VRL formulations + HT (p < .004).

### 3.6. Blood chemistry and histological analysis

As shown in **Figure S4,** analysis of blood 24 h or 7 d post administration of ThermoVRL + HT, NTSL-VRL + HT, or free-VRL +HT did not reveal statistically significant differences in hematocrit (HCT) or hemoglobin concentrations (HGB) compared to untreated and tumor bearing mice. The red blood cell count (RBC) of mice was found to be reduced 7 d post administration of NTSL-VRL + HT (p < .02). Platelet counts (PLT) as well as procalcitonin levels (PCT) were increased in all treatment groups 7 d post administration compared to untreated, tumor bearing control mice (p < .008). No statistically significant differences in alkaline phosphatase (ALP), alanine aminotransferase (ALT), or blood urea nitrogen (BUN) values were detected (p > .25). In conclusion, no noticeable hematological toxicity or changes in liver and kidney parameters were measured at a dose of 15 mg/kg. It is important to note that these measurements were performed after a one-time treatment regimen, whereas animals were treated once per week for a total of three weeks in the efficacy studies.

Liver toxicity was evaluated 24 h and 7 days post administration of free-VRL, NTSL-VRL, or ThermoVRL, all in combination with mild HT. No tissue damage was observed for any of the treatments at either timepoint after a single treatment at a dose of 15 mg/kg (via H&E, TUNEL, and PAS staining). To determine VRL delivery and treatment efficacy, tumor sections were stained for the cell proliferation marker Ki-67. As shown in **Figure 6,** no difference in %Ki-67 positive cells was found 24 h post administration in animals treated with free-VRL +HT or NTSL-VRL +HT (95 ± 2 % and 94 ±3 % respectively). However, the number of proliferating cells was significantly reduced when animals were treated with ThermoVRL + HT (82 ±9 %Ki-67 positive cells; p < .04). No difference in Ki-67 positive cells between the different treatment groups + HT was found seven days post administration.

Ki-67 is employed here as an indirect marker of drug effect [39]. VRL’s anti-mitotic mechanism of action inhibits cell proliferation, thus leading to a reduction in Ki-67 expression. However, for liposomal formulations of VRL the drug must be released in order to exert its effects. NTSL-VRL and ThermoVRL delivered comparable total amounts of VRL (encapsulated and free) to the tumor, yet 24 h post administration cell proliferation was significantly reduced in tumors treated with ThermoVRL compared to NTSL-VRL. This does suggest that intravascular drug release from thermosensitive liposomes results in a more rapid reduction in tumor cell proliferation compared to VRL delivery via NTSL-VRL.

## 4. Discussion

### Influence of D/L ratio on the treatment efficacy of ThermoVRL

Effective intravascular heat-triggered drug delivery relies on stable retention of drug in liposomes at body temperature and rapid burst release once the nanoparticles have reached the heated target area [27]. Release of drug must then be followed by extravasation of the small molecules from the tumor vasculature into the surrounding tumor tissue [24]. Tumor transit times are reported to only be in the range of a few seconds, which is why rapid drug release is key to successful deposition at the target site [40]. Another factor playing an important role is the temperature distribution across the target area. Localized heating of tumors commonly results in a heterogenous temperature distribution across the target volume, where specifically the tumor periphery is heated to temperatures that are at the lower end of the mild hyperthermia range (~39°C) [41,42]. Both, the transit time, and the temperature distribution determine the spatial and temporal window of triggered release from thermosensitive liposomes. Thus, it is not only the release behaviour at ideal mild hyperthermic temperatures (~ 42 °C) that determines the successful release and deposition of the drug from a thermosensitive liposome at the target site, but also the release characteristics at intermediate temperatures (37-40 °C).

In a recently published study, we reported that a decrease in the amount of drug loaded resulted in an increase in the *in vitro* stability of a thermosensitive liposome formulation of VRL [25]. The key difference between the two lead candidate formulations tested in the current study was found to be in their *in vitro* drug release behaviour at intermediate temneratures (39-40 °C): linosomes loaded at the lowest drug-to-lipid ratio exhibited increased stability at 39 °C while achieving rapid release once heated to temperatures above 40 °C. For the reasons mentioned above, we evaluated whether this difference in *in vitro* stability would translate to an impact in *in vivo* efficacy. However, while our study did not find any statistically significant differences in efficacy for mice treated with thermosensitive liposomes loaded with VRL at different D/L ratios, it did appear that thermosensitive liposomes loaded with less drug (i.e., having increased stability) resulted in an increased survival benefit. Thus, one may infer that the increased *in vitro* stability of the liposome formulation with the lower drug loading level, at intermediate temperatures, may be beneficial in preventing premature drug release prior to reaching the heated target site.

### The effect of localized mild hyperthermia on treatment efficacy

Localized mild hyperthermia is able to activate and boost a local anti-tumor immune response that, by itself, has been shown to result in tumor growth inhibition [43]. However, to evaluate our ThermoVRL formulation in the context of RMS we employed a commonly used ectopic xenograft mouse model which requires the use of severely immunocompromised mice. The lack of functional B and T lymphocytes in this strain of mice prohibits the anti-tumor immune effects stimulated by mild hyperthermia treatment. Thus, aside from triggering drug release from thermosensitive liposomes, any beneficial effects on improving tumor growth inhibition are solely based on local physiological changes (e.g., changes in blood flow and vascular permeability) [26,44], Accordingly, we did not observe any effects of mild HT alone on tumor growth or animal survival.

Treatment efficacy as well as tumor VRL concentrations following free-VRL treatment were not affected by the addition of mild HT. By contrast, previous studies from our lab found 2- to 3-fold increases in bulk tumor accumulation of free drug (cisplatin, doxorubicin, and alvespimycin) using the same heating protocol that was employed in these studies (25 min at 42 °C) [26,27], Differences in tumor models (e.g., degree of vascularization and blood perfusion) as well as physico-chemical properties of the drugs (e.g., logP, membrane permeability, or protein binding) could in part explain the observed differences. VRL has been shown to readily distribute and penetrate into tissues compared to other commonly used chemotherapy drugs [45], Thus, the physiological changes that are known to be stimulated by mild HT (e.g., increased blood flow and vascular permeability) may result in a less dramatic effect on the tumor distribution of VRL relative to other drugs.

### Localized mild hyperthermia influences drug delivery of traditional liposomes

Adding localized heating to NTSL-VRL did increase VRL tumor concentrations (5-fold increase albeit not statistically significant). As discussed above, heating did not affect the distribution of free drug thus, this increase must either be due to an increase in accumulation of NTSL-VRL liposomes or due to an increase in drug release from NTSL-VRL that was facilitated by the addition of heat. However, *in vitro* release experiments revealed no significant increase in drug release at the elevated temperature over the applied heating duration. Hence, it is likely that the observed increase in tumor VRL concentration is the result of a heat-stimulated enhancement in liposome accumulation.

Accordingly, the addition of mild HT to treatment with NTSL-VRL did not significantly improve treatment efficacy. The methods employed in this study did not distinguish between the deposition of encapsulated and free drug in the tissue/tumor. Encapsulated drug behaves like a pro-drug: drug release from the liposome is required prior to exerting its cytotoxic activity. Thus, increased liposome accumulation facilitated by the addition of mild HT does not necessarily equate to an increase in bioavailable free drug molecules.

### Delivery of VRL via thermosensitive liposomes

In contrast, the addition of mild HT dramatically improved the treatment efficacy associated with ThermoVRL. *In vitro* drug release experiments previously conducted in our lab demonstrated complete VRL release from the ThermoVRL formulation within seconds of heating to mild hyperthermic temperatures [25]. Maximum release of 95 % was reached after only ~30 s at 42 °C. Additionally, other groups have confirmed rapid release in *in vivo* experiments using a comparable formulation (i.e., same lipid composition): rapid burst release occurs once the thermosensitive liposomes enter the heated blood vessels followed by distribution of drug into the surrounding tissue [46]. Thus, it can be argued that the increase in tumor VRL concentration following the addition of mild HT to the administration of ThermoVRL is due to local burst release of drug within the tumor blood vessels followed by penetration into the surrounding tissue.

Ten Hagen et al. recently demonstrated that there are two key parameters that determine the maximum amount of drug that can be delivered to the target site via intravascular triggered drug delivery systems: the drug’s release kinetics and its extraction from the blood vessels into the surrounding tissue [47], Rapid release and rapid uptake into the tissue are crucial. As discussed earlier, the current first-line chemotherapy regimen for the treatment of RMS remains a combination of vincristine (VCR), actinomycin D, and cyclophosphamide. At first glance, it may appear that VCR would be an ideal candidate for encapsulation and delivery via thermosensitive liposomes. Particularly since liposomal encapsulation of VCR is well understood and, more importantly, already available as an approved therapeutic since 2012 (i.e., Marqibo^®^) [48,49]. However, VRL is more lipophilic and thus membrane permeable. It provides increased cellular and tissue uptake compared to any other compound in the vinca alkaloid family [19,50]. From a formulation standpoint, these properties render VRL the superior drug candidate for intravascular triggered drug delivery. And clinically, as discussed earlier, VRL has shown promising activity in the treatment of RMS [15,16].

Liposomal encapsulation of VRL has previously been pursued and achieved by several research groups as well as the Taiwan Liposome Company (TLC178) [19–21,51–54]. The rationale for liposome encapsulation of VRL is commonly based on a) the well-known fact, that liposomal encapsulation can result in improvements in the drug’s therapeutic index [26] and b) the cell-cycle dependence of VRL’s mechanism of action. As a vinca alkaloid, VRL mainly functions as a mitotic spindle inhibitor leading to metaphase arrest [50]. Thus, it has been postulated that prolonged cell exposure is advantageous and required to increase the exposure of as many cells as possible to VRL during the sensitive phase of the cell cycle [53]. It has been shown that increasing the duration of exposure to VRL improves tumor response. As well, it has been found that VRL’s activity is not only exerted during mitosis, VRL also targets cells in the Gi phase. This in turn may explain VRL’s activity on relatively slow growing tumors with low mitotic indices [55,56]. Ultimately, prolonged tumor cell exposure to VRL is desirable, but more importantly, the delivery approach should focus on tumor specificity including the total amount and distribution of bioavailable drug that reaches the target site. Despite the significantly longer half-lives for VRL delivered via traditional liposomes compared to delivery via thermosensitive liposomes, both delivery vehicles deposited similar amounts of drug to the tumor (measured as AUC_20-360 min,tumor_)· However, it is the amount and distribution of bioavailable free drug that determines treatment efficacy. For thermosensitive liposomes, this is achieved using intravascular triggered drug release [57],

### Drug accumulation at the tumor

Bulk tumor drug accumulation measurements performed at specific timepoints only offer a snapshot into the dynamic process of drug delivery to the tumor. The accumulation process at the tumor continues for as long as there is drug present within tumor blood vessels and stands in constant balance with drug being cleared from the tumor. Ideally, real-time continuous measurements are performed to evaluate this process. However, such measurements commonly rely on the use of contrast agents as drug molecule surrogates or fluorescent drug molecules [58]. To capture the accumulation as well as washout of VRL at the tumor, tumor tissues at various timepoints were excised and drug concentrations after tissue homogenization were determined. Analyzing a single timepoint (e.g., immediately once heating is completed) does not offer the full picture. Particularly, when comparing different drug delivery approaches with such different pharmacokinetics and drug release mechanisms as free drug, traditional liposomes, or intravascular releasing thermosensitive liposomes. Most commonly, tumor drug accumulation following thermosensitive liposome administration is assessed immediately post heating [27,36,59,60]. Here, immediately post heating, VRL concentrations in the tumor are significantly higher when administered as ThermoVRL compared to the administration as free drug as well as a non-thermosensitive liposome formulation. However, upon analyzing several other timepoints, it becomes apparent that the total amount of drug delivered to the tumor is similar between the two liposomal delivery systems. Thus, these results demonstrate the need to analyze tumor drug accumulation at various timepoints tailored to the delivery systems as well as the drug of interest when determining target drug depositions. Particularly since these results are of direct relevance to treatment planning and can suggest and inform on the need and timing of multiple heating/administration cycles [60]. To the knowledge of the authors, this data set represents the first complete pharmacokinetic/tumor distribution analysis of a thermosensitive liposomal formulation in combination with laser-mediated localized mild hyperthermia that does not encapsulate doxorubicin. Previous, studies by other research groups have been heavily focused on doxorubicin as it remains to be the most studied thermosensitive liposomal drug candidate. Additionally, as previously demonstrated by Ramajayam et al., the heating technique significantly affects the pharmacokinetics and tumor accumulation of drugs delivered via thermosensitive liposomes [61]. Particularly, untargeted large volume hyperthermia commonly achieved via water bath heating (remaining the most popular heating method in combination with TSLs [36,60,62]) leads to non-specific drug release and thus increased drug clearance and reduced tumor accumulation. This further strengthens the importance and relevance of the data obtained in the studies presented here where a localized, laser-mediated tumor heating approach was employed [63].

### *In vivo* performance of NTSL-VRL and the EPR effect

Even after several decades of liposomal drug delivery research and the approval of various liposomal formulations, translating successful findings from animal studies into the clinic remains challenging. The most prominent example is the gold-standard formulation Doxil^®^/Caelyx^®^ (i.e., pegylated liposomal doxorubicin). While pegylated liposomal doxorubicin has shown impressive improvements in treatment efficacy across various different animal models this has not been found to translate into comparable benefits in the clinic [64]. To date, various reasons have been identified that contribute to this discrepancy [65]. Among others, two key factors are a heavy reliance on passive tumor targeting via the EPR effect and limited drug release from the carrier at the target site. A significant heterogeneity in the EPR effect has been observed in humans [23]. The EPR effect is significantly more common and pronounced in animal tumor models, which is largely due to differences in the tumor growth rate between tumors in rodent models versus humans [66]. Additionally, the tumor model employed in these studies was found to be well vascularized, which further promotes passive targeting with traditional liposomes. While the traditional NTSL liposome formulation of VRL produced a remarkable improvement in treatment efficacy over the administration of free drug, it does raise the question of clinical translatability. This treatment approach continues to rely on passive targeting via the EPR effect and thus faces the same clinical translation challenges that many nanomedicine-based approaches have failed to overcome in the past. And while we acknowledge that the clinical translation of the multimodal treatment approach presented here comes with its own set of challenges, it does offer a promising strategy to overcome some of the hurdles that are known to be associated with liposomal delivery. It is also important to note that similar levels of bulk tumor accumulation of drug do not necessarily translate into equivalent treatment efficacy. Drug bioavailability as well as drug distribution across the target volume play a key role. This is significantly impacted by limited drug release at the target site, which is common in the case of delivery via traditional liposomes [67]. It is important to note, that the non-thermosensitive liposome formulation used here compares favourably to the aforementioned formulation Doxil/Caelyx. While NTSL-VRL released approximately 30 % of its encapsulated content over the course of seven days, Eetezadi et al. reported virtually no drug release from Caelyx under similar *in vitro* experimental conditions [68]. Externally triggering drug release via localized heating does assure rapid as well as efficient drug release from thermosensitive liposomes and thus increases the amount and distribution of bioavailable drug delivered to the tumor.

### Achieving tumor specific delivery of VRL

One of the key findings of this study is that the delivery of VRL via thermosensitive liposomes significantly improved drug partitioning into the tumor **(Table 2).** As expected, free drug was found to partition into all tissues evaluated equally well (as shown in **Figure 8).** However, encapsulation of VRL into liposomal carriers affected the partitioning behaviour of the encapsulated drug: VRL delivery in non-thermosensitive liposomes significantly reduced tissue as well as tumor partitioning. In contrast, VRL administration via thermosensitive liposomes in combination with localized mild HT at the tumor site showed a significant increase in tumor partitioning of VRL when compared to healthy tissues as well as in comparison to administration via non-thermosensitive liposomes. Hence, the administration of VRL as a thermosensitive liposome formulation provided improved tumor specificity compared to not only administration as free drug but more importantly as a traditional liposome formulation **(Figure 8).**

### Conclusion

Childhood cancer survivors face a significantly longer life expectancy than adult cancer patients [69]. However, RMS survivors are at a 5-fold increased risk of developing a second malignant neoplasm at some point in their life [70]. While ultimately a plethora of factors contribute to this increased risk, off-target delivery of chemotherapy drugs does play an important role [71], Thus, adverse effects due to off-target distribution of chemotherapy drugs are to be avoided at all costs, particularly in children. None of the current standard chemotherapy regimens in the treatment of RMS include targeted therapeutics (vincristine, actinomycin D, and cyclophosphamide in North America and ifosfamide, vincristine, and actinomycin D in Europe). The delivery approach presented here is able to confer targetability to an untargeted drug molecule. We demonstrate the advantage of a targeted delivery approach over the administration of free drug or untargeted, passive drug carriers. This delivery platform increased the amount of bioavailable drug delivered specifically to the tumor while reducing off-target drug deposition. And more importantly, this platform and approach are adaptable and could in the future be expanded to various other commonly used chemotherapy drugs.

## Supporting information

Supplementary Informaton

## 5. Acknowledgements

These studies were supported by a CIHR project grant to C.A. MR holds a Centre for Pharmaceutical Oncology scholarship. The authors acknowledge the use of equipment in the Centre for Pharmaceutical Oncology (CPO) at the University of Toronto and the STTARR Innovation Centre (University Health Network) as well as Professor Heiko Heerklotz’s helpful input on the manuscript.

## 7. Abbreviations

ALP: alkaline phosphatase
ALT: alanine aminotransferase
ATR: ataxia telangiectasia and Rad3-related protein
AUC: area under the curve
BSA: bovine serum albumin
BUN: blood urea nitrogen
CBC: complete blood count
CL: clearance
COG: Children’s Oncology Group
D/L: drug-to-lipid
DLS: dynamic light scattering
DPPC: 1,2-Dipalmitoyl-sn-glycero-3-phosphocholine
DSC: differential scanning calorimetry
DSPC: 1,2-Distearoyl-sn-glycero-3-phosphocholine
EPR: enhanced permeability and retention
FBS: fetal bovine serum
H/E: hematoxylin and eosin
HBS: HEPES-buffered saline
HCT: hematocrit
HDAC: histone deacetylase
HEPES: 4-(2-hydroxyethyl)-1-piperazineethanesulfonic acid
HGB: hemoglobin concentrations
HPLC-MS: high performance liquid chromatography-mass spectrometry
HT: hyperthermia
i.v.: intravenous
IC_50_: half maximal inhibitory concentration
ITS: insulin-transferrin-selenium
K_p,tissue:blood_: tissue and tumor partitioning coefficient
lyso-SPC: 1-stearoyl-2-lyso-sn-glycero-3-phosphocholine
MeOH: methanol
MPS: mononuclear phagocyte system
MST: median survival time
MWCO: molecular weight cut-off
Na_8_SOS: sodium sucrose octasulfate
NaOH: sodium hydroxide
NTSL-VRL: non-thermosensitive liposome formulation encapsulating VRL
OCT: optimal cutting temperature
P/S: penicillin and streptomycin
PARP: poly ADP ribose polymerase
PAS: Periodic acid-Schiff
PBS: phosphate buffered saline
PCT: procalcitonin levels
PEG_2k_-DSPE: N-(carbonyl-methoxypolyethylenglycol 2000)-1,2-distearoyl-sn-glycero-3-phosphoethanolamine
PES: polyethersulfone
PLT: platelet counts
RBC: red blood cell count
RMS: rhabdomyosarcoma
SCID: severe combined immunodeficiency
SD: standard deviation
STS: soft tissue sarcoma
TEA_8_SOS: triethylammonium sucrose octasulfate
ThermoVRL: thermosensitive liposome formulation encapsulating VRL
T_m_: melting phase transition temperature
TUNEL: terminal deoxynucleotidyl transferase dUTP nick end labeling
UHN: University Health Network
VAC: vincristine, actinomycin D, cyclophosphamide
VB: vinblastine sulfate
VCR: vincristine
VRL: vinorelbine tartrate

## References

[1] S.C. Government of Canada, Childhood cancer incidence and mortality in Canada, (2015). https://www.statcan.gc.ca/pub/82-624-x/2015001/article/14213-eng.htm#a1 (accessed November 26, 2017).

[2] E. Butler, K. Ludwig, H.L. Pacenta, L.J. Klesse, T.C. Watt, T.W. Laetsch, Recent progress in the treatment of cancer in children, CA. Cancer J. Clin. 71 (2021) 315–332. https://doi.org/10.3322/caac.21665.

[3] M Fatih Okcu, John Hicks, Rhabdomyosarcoma in childhood, adolescence, and adulthood: Treatment, (2022). uptodate.com (accessed March 2, 2022).

[4] Survival Rates for Rhabdomyosarcoma by Risk Group, (n.d.). https://www.cancer.org/cancer/rhabdomyosarcoma/detection-diagnosis-staging/staging-survival-rates.html (accessed May 8, 2022).

[5] S. Mazzoleni, G. Bisogno, A. Garaventa, G. Cecchetto, A. Ferrari, G. Sotti, A. Donfrancesco, E. Madon, L. Casula, M. Carli, Associazione Italiana di Ematologia e Oncologia Pediatrica Soft Tissue Sarcoma Committee, Outcomes and prognostic factors after recurrence in children and adolescents with nonmetastatic rhabdomyosarcoma, Cancer. 104 (2005) 183–190. https://doi.org/10.1002/cncr.21138.

[6] C.M. Heske, L. Mascarenhas, Relapsed Rhabdomyosarcoma, J. Clin. Med. 10 (2021) 804 https://doi.org/10.3390/jcm10040804.

[7] C. Chen, H. Dorado Garcia, M. Scheer, A.G. Henssen, Current and Future Treatment Strategies for Rhabdomyosarcoma, Front. Oncol. 9 (2019). https://www.frontiersin.org/article/10.3389/fonc.2019.01458 (accessed May 8, 2022).

[8] A.E.M. van Erp, Y.M.H. Versleijen-Jonkers, W.T.A. van der Graaf, E.D.G. Fleuren, Targeted Therapy-based Combination Treatment in Rhabdomyosarcoma, Mol. Cancer Ther. 17 (2018) 1365–1380. https://doi.org/10.1158/1535-7163.MCT-17-1131.

[9] L. Mascarenhas, Y.-Y. Chi, P. Hingorani, J.R. Anderson, E.R. Lyden, D.A. Rodeberg, DJ. Indelicato, S.C. Kao, R. Dasgupta, S.L. Spunt, W.H. Meyer, D.S. Hawkins, Randomized Phase II Trial of Bevacizumab or Temsirolimus in Combination With Chemotherapy for First Relapse Rhabdomyosarcoma: A Report From the Children’s Oncology Group, J. Clin. Oncol. 37 (2019) 2866–2874 https://doi.org/10.1200/JCO.19.00576.

[10] S. Miwa, N. Yamamoto, K. Hayashi, A. Takeuchi, K. Igarashi, H. Tsuchiya, Recent Advances and Challenges in the Treatment of Rhabdomyosarcoma, Cancers. 12 (2020) 1758 https://doi.org/10.3390/cancers12071758.

[11] K.C. Oeffinger, T. Kawashima, D.L. Friedman, N.S. Kadan-Lottick, L.L. Robison, Chronic Health Conditions in Adult Survivors of Childhood Cancer, N Engl J Med. (2006) 11

[12] J.A. Punyko, A.C. Mertens, J.G. Gurney, Y. Yasui, S.S. Donaldson, D.A. Rodeberg, R.B. Raney, M. Stovall, C.A. Sklar, L.L. Robison, K.S. Baker, Long-term medical effects of childhood and adolescent rhabdomyosarcoma: A report from the childhood cancer survivor study, Pediatr. Blood Cancer. 44 (2005) 643–653. https://doi.org/10.1002/pbc.20310.

[13] A. Ruggiero, R. Skinner, W.Z. Khaled Zekri, Editorial: Adverse and Toxic Effects of Childhood Cancer Treatments, Front. Oncol. 11 (2021). https://www.frontiersin.org/article/10.3389/fonc.2021.795664 (accessed March 2, 2022).

[14] S. Hettmer, Z. Li, A.N. Billin, F.G. Barr, D.D.W. Cornelison, A.R. Ehrlich, D.C. Guttridge, A. Hayes-Jordan, L.J. Helman, P.J. Houghton, J. Khan, D.M. Langenau, C.M. Linardic, R. Pal, T.A. Partridge, G.K. Pavlath, R. Rota, B.W. Schäfer, J. Shipley, B. Stillman, L.H. Wexler, A.J. Wagers, C. Keller, Rhabdomyosarcoma: Current Challenges and Their Implications for Developing Therapies, Cold Spring Harb. Perspect. Med. 4 (2014) a025650 https://doi.org/10.1101/cshperspect.a025650.

[15] J.F. Kuttesch, M.D. Krailo, T. Madden, M. Johansen, A. Bleyer, Phase II Evaluation of Intravenous Vinorelbine (Navelbine) in Recurrent or Refractory Pediatric Malignancies: A Children’s Oncology Group Study, Pediatr. Blood Cancer. 53 (2009) 590–593. https://doi.org/10.1002/pbc.22133.

[16] M. Casanova, A. Ferrari, F. Spreafico, M. Terenziani, M. Massimino, R. Luksch, G. Cefalo, D. Polastri, I. Marcon, F.F. Bellani, Vinorelbine in previously treated advanced childhood sarcomas, Cancer. 94 (2002) 3263–3268. https://doi.org/10.1002/cncr.10600.

[17] G. Bisogno, G.L. De Salvo, C. Bergeron, S. Gallego Melcón, J.H. Merks, A. Kelsey, H. Martelli, V. Minard-Colin, D. Orbach, H. Glosli, J. Chisholm, M. Casanova, I. Zanetti, C. Devalck, M. Ben-Arush, P. Mudry, S. Ferman, M. Jenney, A. Ferrari, Vinorelbine and continuous low-dose cyclophosphamide as maintenance chemotherapy in patients with high-risk rhabdomyosarcoma (RMS 2005): a multicentre, open-label, randomised, phase 3 trial, Lancet Oncol. 20 (2019) 1566–1575 https://doi.org/10.1016/S1470-2045(19)30617-5.

[18] A. Ferrari, P. Gasparini, M. Casanova, A home run for rhabdomyosarcoma after 30 years: What now?, Tumori J. 106 (2020) 5–11. https://doi.org/10.1177/0300891619888021.

[19] S.C. Semple, R. Leone, J. Wang, E.C. Leng, S.K. Klimuk, M.L. Eisenhardt, Z.-N. Yuan, K. Edwards, N. Maurer, M.J. Hope, P.R. Cullis, Q.-F. Ahkong, Optimization and characterization of a sphingomyelin/cholesterol liposome formulation of vinorelbine with promising antitumor activity, J. Pharm. Sci. 94 (2005) 1024–1038. https://doi.org/10.1002/jps.20332.

[20] D.C. Drummond, C.O. Noble, Z. Guo, M.E. Hayes, J.W. Park, C.-J. Ou, Y.-L. Tseng, K. Hong, D.B. Kirpotin, Improved Pharmacokinetics and Efficacy of a Highly Stable Nanoliposomal Vinorelbine, J. Pharmacol. Exp. Ther. 328 (2009) 321–330. https://doi.org/10.1124/jpet.108.141200.

[21] C. Li, J. Cui, C. Wang, L. Zhang, X. Xiu, Y. Li, N. Wei, Y. Li, L. Zhang, Encapsulation of vinorelbine into cholesterol-polyethylene glycol coated vesicles: drug loading and pharmacokinetic studies, J. Pharm. Pharmacol. 63 (2011) 376–384. https://doi.org/10.1111/j.2042-7158.2010.01227.x.

[22] S.N. Ekdawi, J.M.P. Stewart, M. Dunne, S. Stapleton, N. Mitsakakis, Y.N. Dou, D.A. Jaffray, C. Allen, Spatial and temporal mapping of heterogeneity in liposome uptake and microvascular distribution in an orthotopic tumor xenograft model, J. Controlled Release. 207 (2015) 101–111. https://doi.org/10.1016/j.jconrel.2015.04.006.

[23] H. Lee, A.F. Shields, B.A. Siegel, K.D. Miller, I. Krop, C.X. Ma, P.M. LoRusso, P.N. Munster, K. Campbell, D.F. Gaddy, S.C. Leonard, E. Geretti, S.J. Blocker, D.B. Kirpotin, V. Moyo, T.J. Wickham, B.S. Hendriks, 64Cu-MM-302 Positron Emission Tomography Quantifies Variability of Enhanced Permeability and Retention of Nanoparticles in Relation to Treatment Response in Patients with Metastatic Breast Cancer, Clin. Cancer Res. 23 (2017) 4190–4202. https://doi.org/10.1158/1078-0432.CCR-16-3193.

[24] B. Kneidl, M. Peller, G. Winter, L.H. Lindner, M. Hossann, Thermosensitive liposomal drug delivery systems: state of the art review, Int. J. Nanomedicine. 9 (2014) 4387–4398. https://doi.org/10.2147/IJN.S49297.

[25] M. Regenold, J. Steigenberger, E. Siniscalchi, M. Dunne, L. Casettari, H. Heerklotz, C. Allen, Determining critical parameters that influence in vitro performance characteristics of a thermosensitive liposome formulation of vinorelbine, J. Controlled Release. (2020). https://doi.org/10.1016/j.jconrel.2020.08.059.

[26] Y.N. Dou, J. Zheng, W.D. Foltz, R. Weersink, N. Chaudary, D.A. Jaffray, C. Allen, Heat-activated thermosensitive liposomal cisplatin (HTLC) results in effective growth delay of cervical carcinoma in mice, J. Controlled Release. 178 (2014) 69–78. https://doi.org/10.1016/j.jconrel.2014.01.009.

[27] M. Dunne, B. Epp-Ducharme, A.M. Sofias, M. Regenold, D.N. Dubins, C. Allen, Heat-activated drug delivery increases tumor accumulation of synergistic chemotherapies, J. Controlled Release. (2019). https://doi.org/10.1016/j.jconrel.2019.06.012.

[28] N. Banerjee, H. Fonge, A. Mikhail, R.M. Reilly, R. Bendayan, C. Allen, Estrone-3-Sulphate, a Potential Novel Ligand for Targeting Breast Cancers, PLoS ONE. 8 (2013) e64O69 https://doi.org/10.1371/journal.pone.0064069.

[29] L. Eklund, M. Bry, K. Alitalo, Mouse models for studying angiogenesis and lymphangiogenesis in cancer, Mol. Oncol. 7 (2013) 259–282. https://doi.org/10.1016/j.molonc.2013.02.007.

[30] B.L. Viglianti, M.W. Dewhirst, R.J. Boruta, J.-Y. Park, C. Landon, A.N. Fontanella, J. Guo, A. Manzoor, C. L. Hofmann, G.M. Palmer, Systemic anti-tumour effects of local thermally sensitive liposome therapy, Int. J. Hyperth. Off. J. Eur. Soc. Hyperthermic Oncol. North Am. Hyperth. Group. 30 (2014) 385–392. https://doi.org/10.3109/02656736.2014.944587.

[31] de Smet M., MR-HIFU mediated local drug delivery using temperature-sensitive liposomes, Technische Universiteit Eindhoven, 2013.

[32] D.-S. Yim, S. Choi, Predicting human pharmacokinetics from preclinical data: volume of distribution, Transl. Clin. Pharmacol. 28 (2020) 169 https://doi.org/10.12793/tcp.2020.28.e19.

[33] Y. Li, P. Xu, D. He, B. Xu, J. Tu, Y. Shen, Long-Circulating Thermosensitive Liposomes for the Targeted Drug Delivery of Oxaliplatin, Int. J. Nanomedicine. 15 (2020) 6721–6734. https://doi.org/10.2147/IJN.S250773.

[34] P. Cressey, M. Amrahli, P.-W. So, W. Gedroyc, M. Wright, M. Thanou, Image-guided thermosensitive liposomes for focused ultrasound enhanced co-delivery of carboplatin and SN-38 against triple negative breast cancer in mice, Biomaterials. 271 (2021) 120758 https://doi.org/10.1016/j.biomaterials.2021.120758.

[35] S.M. Park, M.S. Kim, S.-J. Park, E.S. Park, K.-S. Choi, Y. Kim, H.R. Kim, Novel temperature-triggered liposome with high stability: Formulation, in vitro evaluation, and in vivo study combined with high-intensity focused ultrasound (HIFU), J. Controlled Release. 170 (2013) 373–379. https://doi.org/10.1016/j.jconrel.2013.06.003.

[36] T. Tagami, J.P. May, M.J. Ernsting, S.-D. Li, A thermosensitive liposome prepared with a Cu2+ gradient demonstrates improved pharmacokinetics, drug delivery and antitumor efficacy, J. Controlled Release. 161 (2012) 142–149. https://doi.org/10.1016/j.jconrel.2012.03.023.

[37] I. Robieux, R. Sorio, E. Borsatti, R. Cannizzaro, V. Vitali, P. Aita, A. Freschi, E. Galligioni, S. Monfardini, Pharmacokinetics of vinorelbine in patients with liver metastases, Clin. Pharmacol. Ther. 59 (1996) 32–40. https://doi.org/10.1016/S0009-9236(96)90021-1.

[38] M.L. Immordino, F. Dosio, L. Cattel, Stealth liposomes: review of the basic science, rationale, and clinical applications, existing and potential, Int. J. Nanomedicine. 1 (2006) 297–315.

[39] T.O. Nielsen, S.C.Y. Leung, D.L. Rimm, A. Dodson, B. Acs, S. Badve, C. Denkert, M.J. Ellis, S. Fineberg, M. Flowers, H.H. Kreipe, A.-V. Laenkholm, H. Pan, F.M. Penault-Llorca, M.-Y. Polley, R. Salgado, I.E. Smith, T. Sugie, J.M.S. Bartlett, L.M. McShane, M. Dowsett, D.F. Hayes, Assessment of Ki67 in Breast Cancer: Updated Recommendations From the International Ki67 in Breast Cancer Working Group, JNCI J. Natl. Cancer Inst. 113 (2021) 808–819. https://doi.org/10.1093/jnci/djaa201.

[40] C. Burke, M.R. Dreher, A.H. Negussie, A.S. Mikhail, P. Yarmolenko, A. Patel, B. Skilskyj, B.J. Wood, D. Haemmerich, Drug release kinetics of temperature sensitive liposomes measured at high temporal resolution with a millifluidic device, Int. J. Hyperth. Off. J. Eur. Soc. Hyperthermic Oncol. North Am. Hyperth. Group. 34 (2018) 786–794. https://doi.org/10.1080/02656736.2017.1412504.

[41] G.C. van Rhoon, M. Franckena, T.L.M. ten Hagen, A moderate thermal dose is sufficient for effective free and TSL based thermochemotherapy, Adv. Drug Deliv. Rev. (2020). https://doi.org/10.1016/j.addr.2020.03.006.

[42] A. Partanen, P.S. Yarmolenko, A. Viitala, S. Appanaboyina, D. Haemmerich, A. Ranjan, G. Jacobs, D. Woods, J. Enholm, B.J. Wood, M.R. Dreher, Mild hyperthermia with magnetic resonance-guided high-intensity focused ultrasound for applications in drug delivery, Int. J. Hyperthermia. 28 (2012) 320–336. https://doi.org/10.3109/02656736.2012.680173.

[43] S. Toraya-Brown, S. Fiering, Local tumour hyperthermia as immunotherapy for metastatic cancer, Int. J. Hyperthermia. 30 (2014) 531–539. https://doi.org/10.3109/02656736.2014.968640.

[44] M. Dunne, M. Regenold, C. Allen, Hyperthermia can alter tumor physiology and improve chemo-and radio-therapy efficacy, Adv. Drug Deliv. Rev. 163-164 (2020) 98–124. https://doi.org/10.1016/j.addr.2020.07.007.

[45] A. Kyle, A. Minchinton, Classification of anticancer drugs based on tissue penetration using a novel in vitro screening assay, Mol. Cancer Ther. 6 (2007) A213

[46] M.A. Santos, S.-K. Wu, M. Regenold, C. Allen, D.E. Goertz, K. Hynynen, Novel fractionated ultrashort thermal exposures with MRI-guided focused ultrasound for treating tumors with thermosensitive drugs, Sci. Adv. 6 (2020) eaba5684 https://doi.org/10.1126/sciadv.aba5684.

[47] T.L.M. ten Hagen, M.R. Dreher, S. Zalba, A.L.B. Seynhaeve, M. Amin, L. Li, D. Haemmerich, Drug transport kinetics of intravascular triggered drug delivery systems, Commun. Biol. 4 (2021) 920 https://doi.org/10.1038/s42003-021-02428-z.

[48] L.D. Mayer, M.B. Bally, H. Loughrey, D. Masin, P.R. Cullis, Liposomal Vincristine Preparations Which Exhibit Decreased Drug Toxicity and Increased Activity against Murine L1210 and P388 Tumors1, Cancer Res. 50 (1990) 575–579.

[49] J.A. Silverman, S.R. Deitcher, Marqibo^®^ (vincristine sulfate liposome injection) improves the pharmacokinetics and pharmacodynamics of vincristine, Cancer Chemother. Pharmacol. 71 (2013) 555–564. https://doi.org/10.1007/s00280-012-2042-4.

[50] E. Rowinsky, The Vinca Alkaloids, Holl.-Frei Cancer Med. 6th Ed. (2003). http://www.ncbi.nlm.nih.gov/books/NBK12718/ (accessed February 22, 2022).

[51] H. Zhang, Z. Wang, W. Gong, Z. Li, X. Mei, W. Lv, Development and characteristics of temperature-sensitive liposomes for vinorelbine bitartrate, Int. J. Pharm. 414 (2011) 56–62. https://doi.org/10.1016/j.ijpharm.2011.05.013.

[52] C.L. Li, J.X. Cui, C.X. Wang, L. Zhang, Y.H. Li, L. Zhang, X. Xiu, Y.F. Li, N. Wei, Development of pegylated liposomal vinorelbine formulation using “post-insertion” technology, Int. J. Pharm. 391 (2010) 230–236. https://doi.org/10.1016/j.ijpharm.2010.03.004.

[53] I.V. Zhigaltsev, N. Maurer, Q.-F. Akhong, R. Leone, E. Leng, J. Wang, S.C. Semple, P.R. Cullis, Liposome-encapsulated vincristine, vinblastine and vinorelbine: A comparative study of drug loading and retention, J. Controlled Release. 104 (2005) 103–111. https://doi.org/10.1016/j.jconrel.2005.01.010.

[54] Taiwan Liposome Company, TLC178-Taiwan Liposome Company, (n.d.). https://www.tlcbio.com/en-global/pipeline/index/Oncology/TLC178 (accessed February 22, 2022).

[55] J.K. Horton, P.J. Houghton, J.A. Houghton, Relationships between tumor responsiveness, vincristine pharmacokinetics and arrest of mitosis in human tumor xenografts, Biochem. Pharmacol. 37 (1988) 3995–4000. https://doi.org/10.1016/0006-2952(88)90085-8.

[56] A. Kothari, W.N. Hittelman, T.C. Chambers, Cell Cycle-Dependent Mechanisms Underlie Vincristine-Induced Death of Primary Acute Lymphoblastic Leukemia Cells, Cancer Res. 76 (2016) 3553–3561. https://doi.org/10.1158/0008-5472.CAN-15-2104.

[57] T. Lu, D. Haemmerich, H. Liu, A.L.B. Seynhaeve, G.C. van Rhoon, A.B. Houtsmuller, T.L.M. ten Hagen, Externally triggered smart drug delivery system encapsulating idarubicin shows superior kinetics and enhances tumoral drug uptake and response, Theranostics. 11 (2021) 5700–5712. https://doi.org/10.7150/thno.55163.

[58] J.A. Tashjian, M.W. Dewhirst, D. Needham, B.L. Viglianti, Rationale for and measurement of liposomal drug delivery with hyperthermia using non-invasive imaging techniques, Int. J. Hyperth. Off. J. Eur. Soc. Hyperthermic Oncol. North Am. Hyperth. Group. 24 (2008) 79–90. https://doi.org/10.1080/02656730701840147.

[59] C. Kim, Y. Guo, A. Velalopoulou, J. Leisen, A. Motamarry, K. Ramajayam, M. Aryal, D. Haemmerich, C.D. Arvanitis, Closed-loop trans-skull ultrasound hyperthermia leads to improved drug delivery from thermosensitive drugs and promotes changes in vascular transport dynamics in brain tumors, Theranostics. 11 (2021) 7276–7293. https://doi.org/10.7150/thno.54630.

[60] W.T. Al-Jamal, Z.S. Al-Ahmady, K. Kostarelos, Pharmacokinetics & tissue distribution of temperature-sensitive liposomal doxorubicin in tumor-bearing mice triggered with mild hyperthermia, Biomaterials. 33 (2012) 4608–4617. https://doi.org/10.1016/j.biomaterials.2012.03.018.

[61] Untargeted Large Volume Hyperthermia Reduces Tumor Drug Uptake From Thermosensitive Liposomes, IEEE Open J. Eng. Med. Biol. 2 (2021) 187–197. https://doi.org/10.1109/OJEMB.2021.3078843.

[62] W.T. Al-Jamal, K. Kostarelos, Mild hyperthermia accelerates doxorubicin clearance from tumour-extravasated temperature-sensitive liposomes, Nanotheranostics. 6 (2022) 230–242. https://doi.org/10.7150/ntno.61280.

[63] Y.N. Dou, R.A. Weersink, W.D. Foltz, J. Zheng, N. Chaudary, D.A. Jaffray, C. Allen, Custom-designed Laser-based Heating Apparatus for Triggered Release of Cisplatin from Thermosensitive Liposomes with Magnetic Resonance Image Guidance, J. Vis. Exp. JoVE. (2015). https://doi.org/10.3791/53055.

[64] G.H. Petersen, S.K. Alzghari, W. Chee, S.S. Sankari, N.M. La-Beck, Meta-analysis of clinical and preclinical studies comparing the anticancer efficacy of liposomal versus conventional non-liposomal doxorubicin, J. Controlled Release. 232 (2016) 255–264. https://doi.org/10.1016/j.jconrel.2016.04.028.

[65] D. Sun, S. Zhou, W. Gao, What Went Wrong with Anticancer Nanomedicine Design and How to Make It Right, ACS Nano. 14 (2020) 12281–12290. https://doi.org/10.1021/acsnano.9b09713.

[66] T. Lammers, F. Kiessling, W.E. Hennink, G. Storm, Drug targeting to tumors: Principles, pitfalls and (pre-) clinical progress, J. Controlled Release. 161 (2012) 175–187. https://doi.org/10.1016/j.jconrel.2011.09.063.

[67] K.M. Laginha, S. Verwoert, G.J.R. Charrois, T.M. Allen, Determination of Doxorubicin Levels in Whole Tumor and Tumor Nuclei in Murine Breast Cancer Tumors, Clin. Cancer Res. 11 (2005) 6944–6949. https://doi.org/10.1158/1078-0432.CCR-05-0343.

[68] S. Eetezadi, R. De Souza, M. Vythilingam, R. Lessa Cataldi, C. Allen, Effects of Doxorubicin Delivery Systems and Mild Hyperthermia on Tissue Penetration in 3D Cell Culture Models of Ovarian Cancer Residual Disease, Mol. Pharm. 12 (2015) 3973–3985. https://doi.org/10.1021/acs.molpharmaceut.5b00426.

[69] H.Y. Ju, E.-K. Moon, J. Lim, B.K. Park, H.Y. Shin, Y.-J. Won, H.J. Park, Second malignant neoplasms after childhood cancer: A nationwide population-based study in Korea, PLoS ONE. 13 (2018) e0207243 https://doi.org/10.1371/journal.pone.0207243.

[70] N.M. Archer, R.P. Amorim, R. Naves, S. Hettmer, L.R. Diller, K.B. Ribeiro, C. Rodriguez-Galindo, An Increased Risk of Second Malignant Neoplasms After Rhabdomyosarcoma: Population-Based Evidence for a Cancer Predisposition Syndrome?, Pediatr. Blood Cancer. 63 (2016) 196–201. https://doi.org/10.1002/pbc.25678.

[71] H. Zhen, Z. Liu, H. Guan, J. Ma, W. Wang, J. Shen, Z. Miao, F. Zhang, Second Malignant Neoplasms in Patients With Rhabdomyosarcoma, Front. Oncol. 11 (2021). https://www.frontiersin.org/article/10.3389/fonc.2021.757095 (accessed February 24, 2022).

