## Supplementary Informaton for "Triggered Release from Thermosensitive Liposomes Improves Tumor Targeting of Vinorelbine"

**Supplementary Information**

**Triggered Release from Thermosensitive Liposomes Improves Tumor Targeting of Commonly Used Chemotherapy Drug Vinorelbine**

Maximilian Regenold^1^, Kan Kaneko^1^, Xuehan Wang^1^, H. Benson Peng^1^, James C. Evans^1^, Pauric Bannigan^1^, Christine Allen^1,*^

^1^Leslie Dan Faculty of Pharmacy, University of Toronto, Toronto, Ontario, Canada

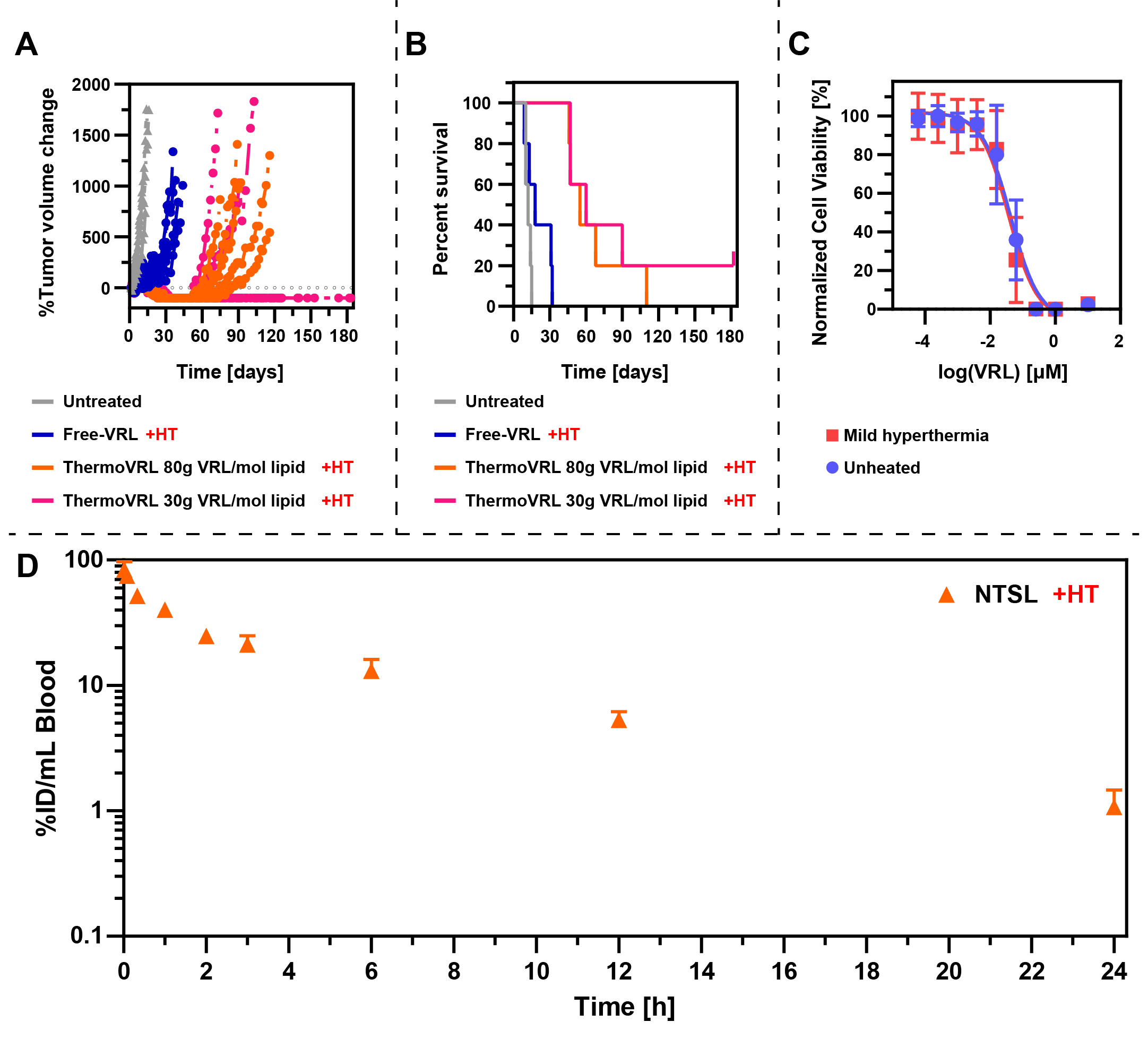

**Figure S1:** A) Rh30 tumor volumes of female SCID mice treated with free vinorelbine (VRL), or thermosensitive liposomes (ThermoVRL) loaded at different drug-to-lipid ratios all in combination with mild hyperthermia (HT; 42.5 °C, 25 min) localized to the tumor site. Treatments were administered at a dose of 15 mg VRL/kg body weight on days 0, 7, and 14 (n = 5). B) Kaplan-Meier survival analysis of tumor bearing female SCID mice treated with VRL administered at 10 mg VRL/kg body weight. C) Evaluation of VRL cytotoxicity on Rh30 cells +/- HT (42 °C for 1 h). Cells were treated with VRL for 1 h, washed and incubated for a total of 72 h. The IC_50_ of VRL in Rh30 cells was found to be 41.4 ± 29.7 nM and 31.5 ± 12.6 nM, without and with the addition of mild hyperthermia, respectively.

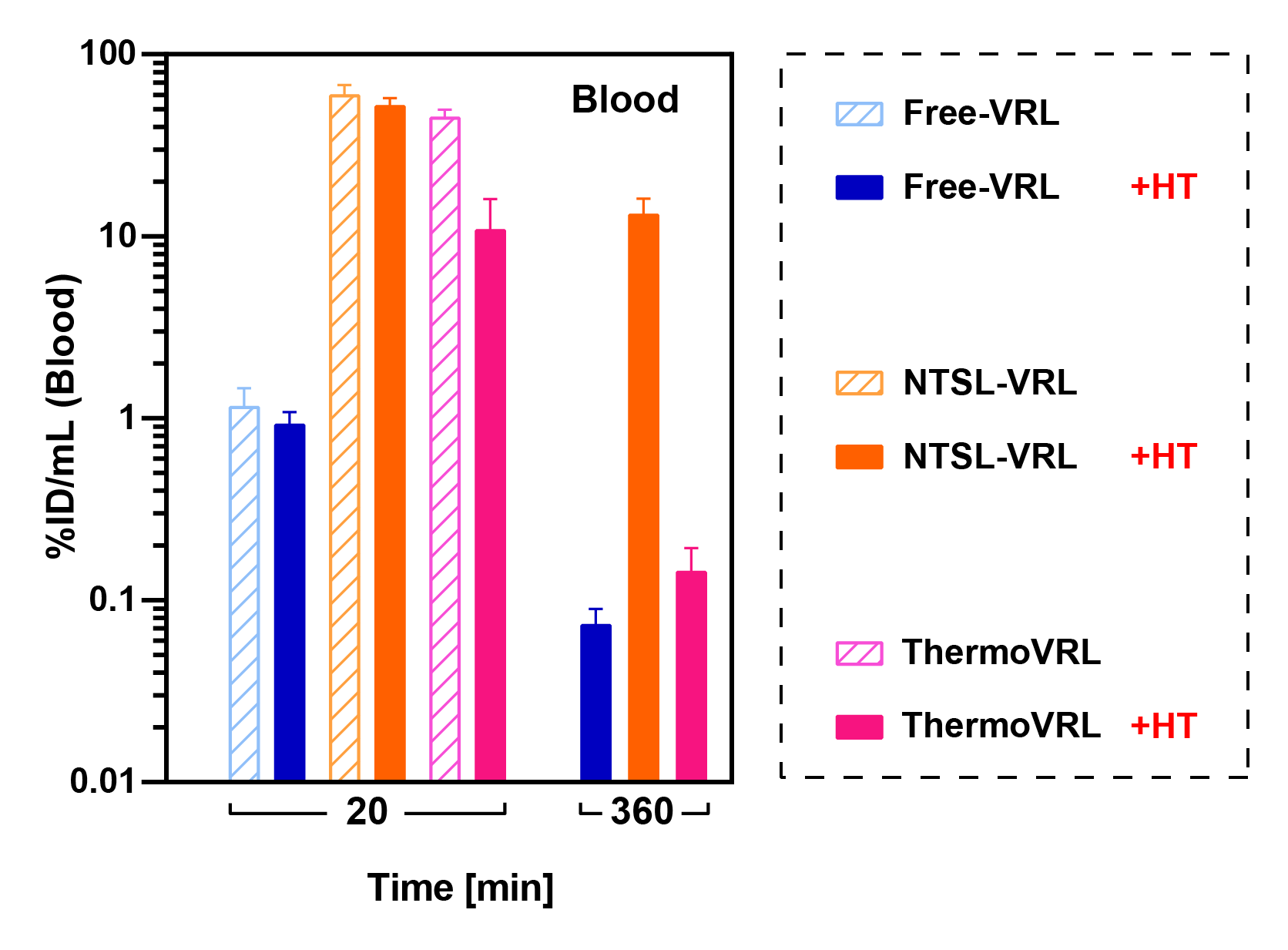

**Figure S2:** VRL concentration in whole blood 20- and 360-min post administration of free vinorelbine (VRL), non-thermosensitive liposomal VRL (NTSL-VRL), or thermosensitive liposomal VRL (ThermoVRL) at 15 mg/kg +/- mild hyperthermia (HT; 42.5 °C, 25 min) localized to the tumor in female SCID mice. Data is presented as mean ± SD (n = 4).

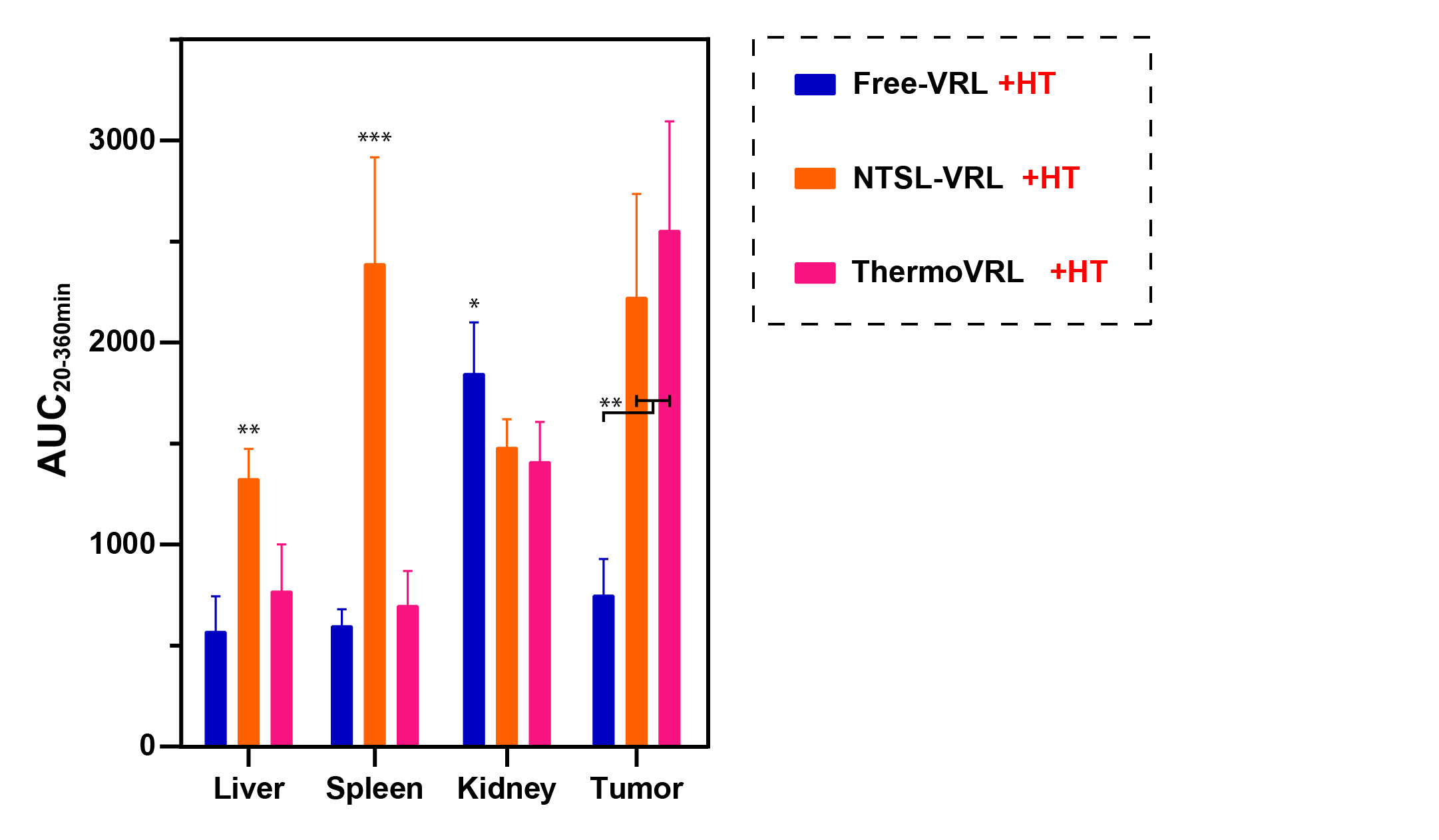

**Figure S3**: AUC_20-360 min_ for vinorelbine (VRL) in different organs and tumors following intravenous administration as free VRL, non-thermosensitive liposomal VRL (NTSL-VRL), and thermosensitive liposomal VRL (ThermoVRL), with each treatment in combination with localized mild hyperthermia (HT) to the tumor (n = 4).

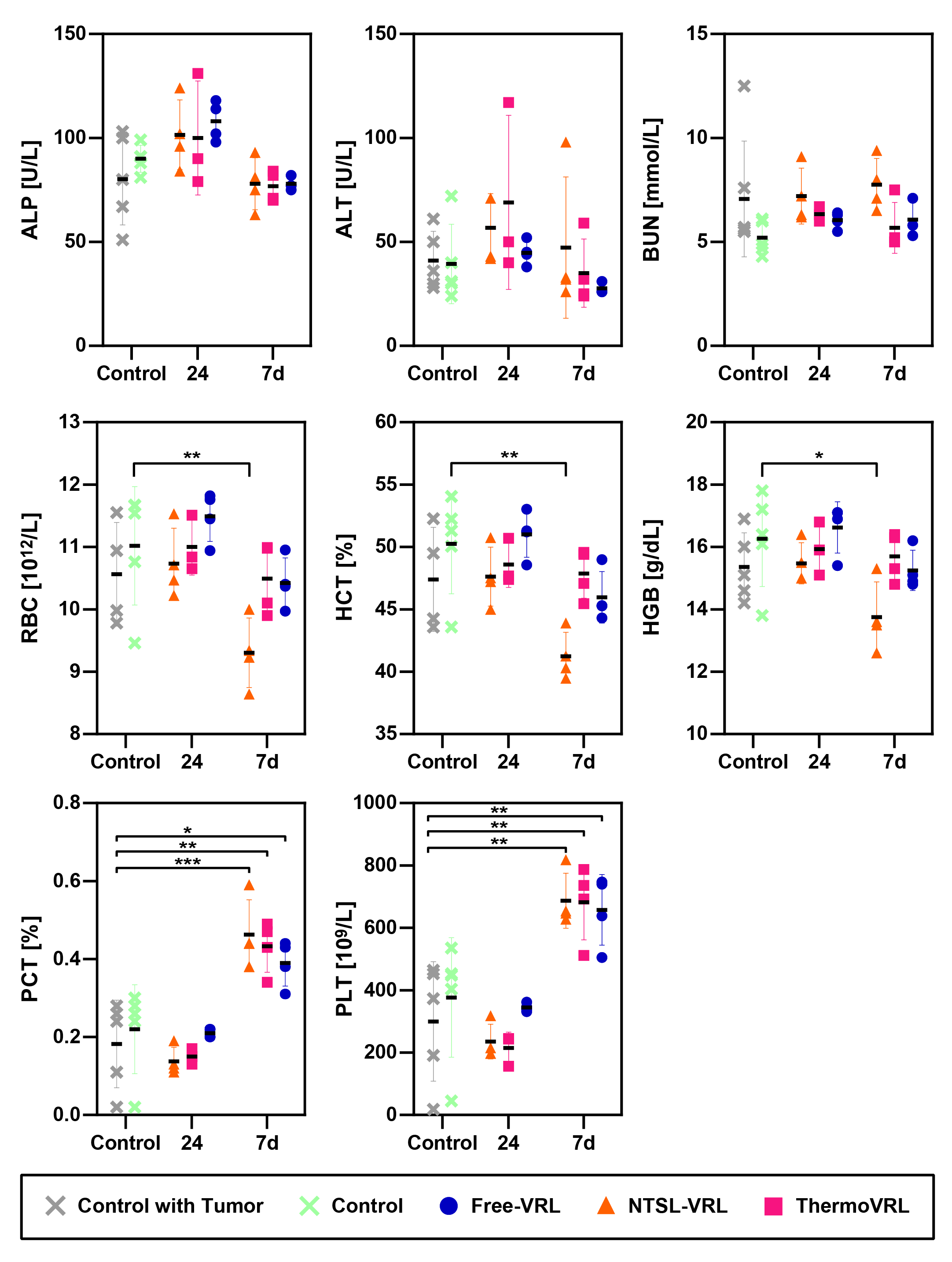

**Figure S4**: Hematological toxicity as well as liver and kidney parameters were analyzed using the VetScan® HM5 Hematology System and the VS2 Chemistry Analyzer. Female SCID mice were treated with vinorelbine (VRL) administered as free-VRL, non-thermosensitive liposomal VRL (NTSL-VRL), or thermosensitive liposomal VRL (ThermoVRL) at 15 mg/kg in combination with mild hyperthermia (HT; 42.5 °C, 25 min) localized to the tumor. Toxicity parameters were analyzed 24 h and 7 days post administration. Tumor bearing (i.e., control with tumor) and non-tumor bearing, untreated animals served as controls. Data is presented as mean ± SD (n ≥ 3).
